# Chrna5 and lynx prototoxins identify acetylcholine super-responder subplate neurons

**DOI:** 10.1101/2022.06.20.496737

**Authors:** Sridevi Venkatesan, Tianhui Chen, Yupeng Liu, Eric E Turner, Shreejoy Tripathy, Evelyn K Lambe

## Abstract

Attention depends on cholinergic excitation of prefrontal neurons but is sensitive to perturbation of α5-containing nicotinic receptors encoded by *Chrna5*. However, *Chrna5*-expressing (Chrna5+) neurons remain enigmatic, despite their potential as a target to improve attention. Here, we generate complex transgenic mice to probe Chrna5+ neurons and their sensitivity to endogenous acetylcholine. Through opto-physiological experiments, we discover that Chrna5+ neurons contain a distinct population of acetylcholine super-responders. Leveraging single-cell transcriptomics, we discover molecular markers conferring subplate identity on this subset. We determine that Chrna5+ super-responders express a unique complement of GPI-anchored lynx prototoxin genes (*Lypd1, Ly6g6e*, and *Lypd6b*), predicting distinct nicotinic receptor regulation. To manipulate lynx regulation of endogenous nicotinic responses, we developed a pharmacological strategy guided by transcriptomic predictions. Overall, we reveal *Chrna5*-Cre mice as a transgenic tool to target the diversity of subplate neurons in adulthood, yielding new molecular strategies to manipulate their cholinergic activation relevant to attention disorders.

## Introduction

Cholinergic modulation of the medial prefrontal cortex (mPFC) is essential for attention and detection of sensory cues^1–4^. Deep-layer pyramidal neurons in the PFC are critically involved in such executive function^5–7^ and are robustly excited by acetylcholine^8,9^ via nicotinic and muscarinic receptor activation^10^. The α5 nicotinic receptor subunit encoded by *Chrna5* is specifically expressed in deep-layer pyramidal neurons^11,12^, forming high-affinity nicotinic receptors in combination with α4 and β2 subunits. Electrophysiological, behavioural, and genetic evidence in both rodents and humans point to an important role of *Chrna5* expression for nicotinic receptor function, attention, and executive function.

Constitutive deletion of *Chrna5* in mice, or knockdown in the adult rat PFC disrupt attention and reduce nicotinic receptor activation by exogenous acetylcholine stimulation in layer 6 neurons^13–5^. Optogenetic experiments in *Chrna5^-/-^* mice show that *Chrna5* is required for rapid onset of postsynaptic cholinergic activation and prevents desensitization of the endogenous cholinergic response during prolonged stimulation^16^. In humans, the non-synonymous rs16969968 (D398N) polymorphism in *Chrna5* is associated with nicotine dependence, schizophrenia and cognitive impairment^17–19^ Nicotinic α4β2α5 receptors with the D398N polymorphism have partial loss of function, attributed to changes in receptor desensitization, calcium permeability, or membrane trafficking^20–22^.

Despite α5 nicotinic receptor involvement in attention and prefrontal cholinergic activation, systematic characterization of *Chrna5-*expressing neurons using modern genetic tools is lacking. Deep-layer neurons include diverse corticothalamic (L6CT), corticocortical, and layer 6b (L6b) populations^23–25^. *Chrna5* is predicted to be expressed in L6CT neurons^8,26^, which are usually identified by their expression of *Syt6*, a conserved L6CT neuronal marker^27,28,29^. *Syt6-* Cre mice have been widely used to characterize prefrontal L6CT neurons and their cholinergic properties^30–32^. However, it is unclear whether these are the same neurons expressing *Chrna5*. Characterization of *Chrna5*-expressing neurons has been limited by the lack of verified antibodies for the α5 subunit that could be used for post-hoc immunostaining. Previous BAC-transgenic mice labeling *Chrna5*-expressing neurons had altered expression of other genes in the tightly linked *Chrna5/a3/b4* gene cluster, limiting their use for functional examination^33^. This issue was circumvented by disrupting the open reading frames of *Chrna3/b4* in the BAC transgene to generate a *Chrna5-Cre* mouse without misexpression artifacts^34^

Here, we create compound transgenic mice to investigate how prefrontal *Chrna5*-expressing (Chrna5+) neurons respond to optogenetic release of endogenous acetylcholine. The fast and strong response prompted a multi-approach examination of Chrna5+ neurons together with a control population of *Syt6*-expressing (Syt6+) neurons, a more traditional molecular marker of deep-layer prefrontal cortex. We demonstrate a large fraction of Chrna5+ neurons have distinctive high-affinity cholinergic responses. Single-cell RNAseq analysis reveals the expression of several subplate markers (*Cplx3, Ctgf*, and *Lpar1*) in this Chrna5+ subset, identifying them as subplate neurons born early in development that are critical for establishing thalamocortical connectivity^35–37^ Chrna5+ subplate neurons show distinct expression pattern of GPI-anchored lynx prototoxins (*Ly6g6e, Lypd1*, and *Lypd6b*) capable of exerting complex modulation of nicotinic receptor properties^38,39^. As predicted from the transcriptomic analysis, our pharmacological manipulations targeting lynx prototoxins successfully alter endogeneous nicotinic responses, revealing cell-type specific lynx regulation of nicotinic receptors.

Recent studies have examined cell-type specific modulation of nicotinic receptors by different lynx prototoxins and the consequences for cortical development and cognition^40–13^. Here, we discovered endogenous lynx modulation of nicotinic properties relevant for attention in subplate neurons expressing *Chrna5*. Our work highlights *Chrna5-Cre* mice as a transgenic tool to target acetylcholine super-responder subplate neurons and identifies strategies to fine-tune their cholinergic activation by manipulating GPI-anchored lynx prototoxins.

## Results

### Chrna5 expression identifies acetylcholine ‘super-responders’

To characterize prefrontal *Chrna5-expressing* (Chrna5+) neurons, we generated triple transgenic *Chrna5*-Cre^/+^Ai14^/+^ChAT-ChR2^/+^ mice (**Fig 1A**) and examined optogenetic cholinergic responses in labeled Chrna5+ neurons. These mice expressed tdTomato in Chrna5+ neurons and channelrhodopsin-2 in cholinergic axons as seen by two-photon imaging (**Fig 1B**). The location of reporter labeled Chrna5+ neurons across the brain in *Chrna5-Cre* mice is specific to layer 6 and found across the entire cortical mantle (**Fig S1**). We recorded endogenous cholinergic responses in labeled Chrna5+ neurons with optophysiology in mPFC slices, using unlabeled pyramidal neurons as a control (**Fig 1C**). Optogenetic Chrna5+ neurons clearly possessed larger-amplitude cholinergic responses with significantly faster onset compared to control neurons (**Fig 1D-F**). The rising slope was significantly larger in Chrna5+ neurons (220 ± 32 pA/s, 24 cells from 4 mice) compared to control (123 ± 22 pA/s, 15 cells; t_(37)_ = 2.19, *P* = 0.02, unpaired t-test; Cohen’s d: 0.72). Similarly, peak current evoked by optogenetic acetylcholine release was greater in Chrna5+ neurons (15 ± 2 pA) compared to control (8 ± 1 pA, t_(37)_ = 2.52, *P* = 0.02; Cohen’s d: 0.83), as well as the area of the cholinergic response (t_(37)_ = 2.06, *P* = 0.046). Intrinsic electrophysiological properties, however, did not differ between Chrna5+ and unlabeled neurons (**Table S1**). Additionally, we determined that Chrna5+ neurons are under presynaptic muscarinic autoinhibitory control, which can be relieved by atropine, resulting in even larger responses (post atropine response: 194 ± 42% of baseline, n = 6 cells). Our application of the nicotinic antagonist DHBE eliminated the optogenetic cholinergic response (post DHBE response: 3 ± 2 % of baseline, n = 4 cells), indicating the relevant nicotinic receptors contain β2 subunits. This characterization of Chrna5+ neurons revealed them to be acetylcholine ‘super-responders’ with stronger and faster onset cholinergic responses distinct from other deep-layer pyramidal neurons. We next examined whether Chrna5+ neurons in other cortical regions are also acetylcholine super-responders. Chrna5+ labeled neurons deep in layer 6 of primary somatosensory cortex (SSp) were also found to show significantly stronger and faster onset nicotinic responses to optogenetic acetylcholine release (**Fig S2)** confirming that *Chrna5*-labeling identifies acetylcholine super-responders in multiple cortical areas.

**Figure 1.**
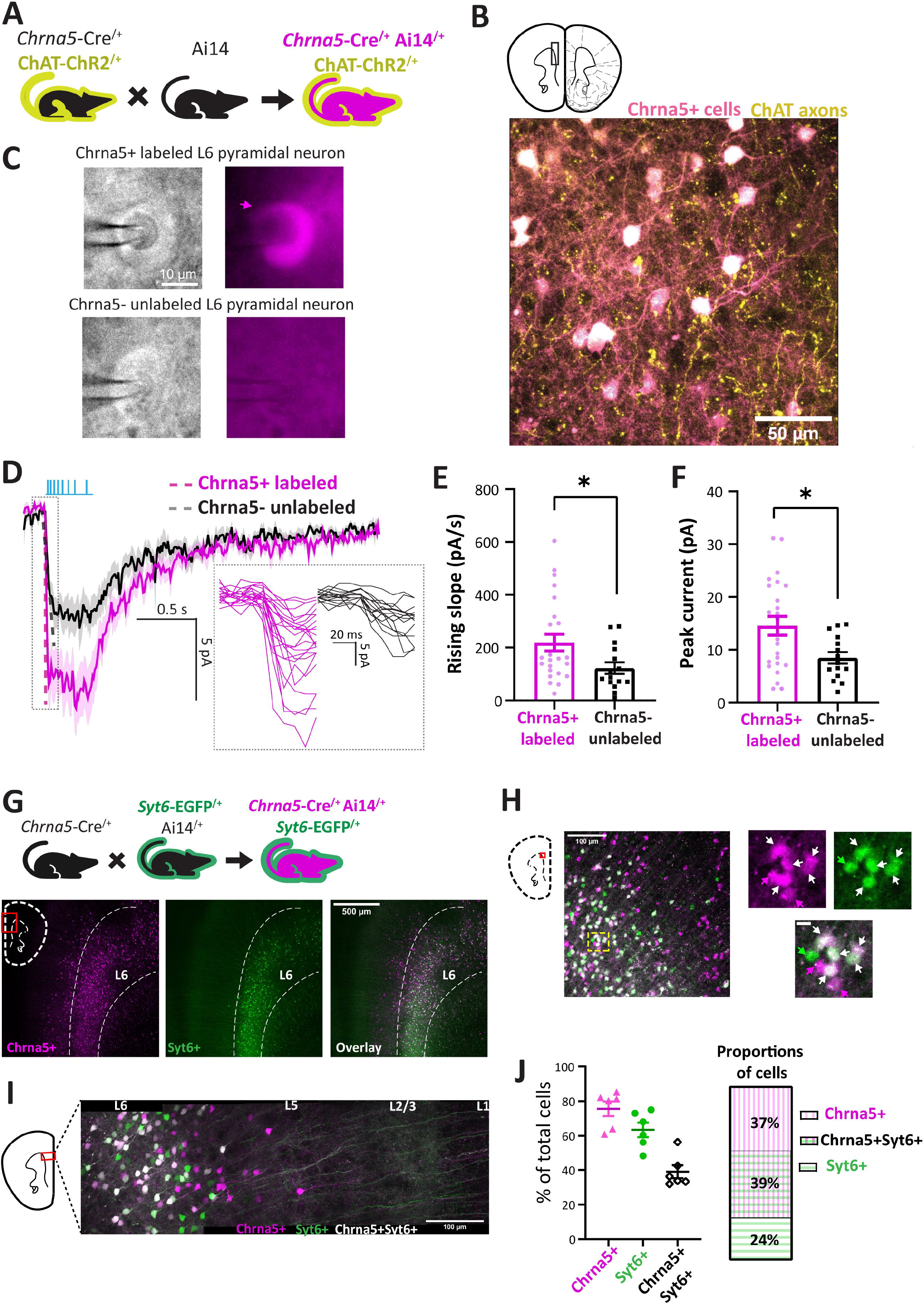
*Chrna5* expression identifies a distinct population of prefrontal neurons with stronger and faster-onset optogenetic cholinergic responses. **A,** Breeding scheme to obtain triple transgenic *Chrna5*-Cre^/+^Ai14^/+^ChAT-ChR2^/+^ mice expressing tdTomato in *Chrna5-* expressing (Chrna5+) neurons and Channelrhodopsin 2 in cholinergic axons. **B**, (*Top*) Schematic of coronal mPFC slice with region of interest, adapted from^89^. (*Bottom*) two photon imaging (3D projection) of tdTomato-labeled Chrna5+ neurons and EYFP-labeled cholinergic axons in layer 6 of mPFC slices. **C**, IRDIC (*left*) and widefield fluorescence (TRITC, *right*) images of tdTomato labeled Chrna5+ and unlabeled layer 6 neurons during whole cell patch clamp electrophysiology. Clearing induced by the pipette is visible. **D**, Average light-evoked endogenous cholinergic response of labeled Chrna5+ vs neighbouring unlabeled Chrna5-neurons. Dotted lines are the slope of the response onset. (*Inset*) Individual responses are zoomed in to show the onset. **E-F**, Bar graph comparing (E) Rising slope and (F) Peak current of endogenous cholinergic responses between labeled Chrna5+ neurons (n = 24 cells) and unlabeled Chrna5-neurons (n = 15 cells, 4 mice). \**P* < 0.05, Unpaired t-test. **G**, (*Top*) Breeding scheme to obtain triple transgenic *Chrna5-* Cre^/+^Ai14^/+^*Syt6*-EGFP^/+^ mice expressing tdTomato in Chrna5+ neurons and EGFP in Syt6+ neurons. (*Bottom*) Confocal imaging in mPFC slices shows Chrna5+ and Syt6+ neurons distributed in layer 6. **H-I,** Confocal (H) and two-photon imaging (I) reveal three populations of neurons: exclusively Chrna5+ neurons which do not express *Syt6*, overlapping Chrna5+Syt6+ neurons which express both markers, and exclusively Syt6+ neurons which do not express *Chrna5*. **J,** (*left*) Graph quantifies the percentage of each cell type with respect to all labeled cells per sample. (*Right*) Average proportions of Chrna5+, Chrna5+Syt6+ and Syt6+ neurons. (See **Figs S1-3** for additional information on distribution of Chrna5+ cells across the cortex, and additional recordings showing stronger and faster opto-cholinergic responses in Chrna5+ neurons in primary somatosensory cortex.)

To further probe the distinct cholinergic responses of prefrontal Chrna5+ neurons, we examined factors differentiating these cells from neurons labeled by *Syt6* expression (Syt6+), which has hereto been the predominant marker of layer 6 corticothalamic neurons used to characterize their function during PFC-dependent tasks^28,30,31,44^ Since the extent of overlap between *Chrna5* and *Syt6* expressing deep-layer pyramidal neuron populations was unclear, we adopted an imaging strategy to visualize the distribution of Chrna5+ and Syt6+ neurons in the PFC and determine the exact proportion of distinctive non-overlapping Chrna5+ neurons. We generated a compound transgenic *Chrna5*-Cre^/+^Ai14^/+^*Syt6*-EGFP^/+^ mouse to simultaneously express tdTomato in Chrna5+ neurons and EGFP in Syt6+ neurons and performed confocal and two-photon imaging of the endogenous fluorescence of these reporters in mPFC brain slices. Chrna5+ and Syt6+ neurons were both present primarily in layer 6, with a few Chrna5+ neurons in layer 5 (**Fig 1G, I**). Fig 1J (left) shows the proportion of all cells expressing tdTomato (*Chrna5*), GFP (*Syt6*), or both, as a percentage of the total number of fluorescent cells in that region. Closer investigation confirmed the existence of a substantial proportion of exclusively Chrna5+ neurons (37% of all labeled cells) which do not express *Syt6*, in addition to overlapping Chrna5+Syt6+ neurons (39%) which express both markers, and exclusively Syt6+ neurons (24%) which do not express *Chrna5* (N = 4 mice, **Fig 1H, J**). Thus, nearly half of all *Chrna5-* expressing neurons are not labeled by *Syt6*-expression and would have been excluded in previous studies using *Syt6*-Cre mice. Furthermore, *Chrna5-expressing* neurons are found even in primary visual cortex, where *Syt6*-labeling is greatly reduced^31,45^ **(Fig S3)**.

### Distinct Chrna5+ neurons with highly-resilient nicotinic receptor responses

To elucidate the distinct subset of Chrna5+ neurons not found with Syt6-labeling approaches, we measured acetylcholine-evoked signals in multiple neurons simultaneously with *ex vivo* GCaMP6s Ca^2+^ imaging. We generated transgenic mice expressing GCaMP6s in either Chrna5+ (*Chrna5*-Cre^/+^/Ai96^/+^) or Syt6+ (*Syt6*-Cre^/+^/Ai96^/+^) neurons and performed two-photon imaging of mPFC layer 6 (**Fig 2A**). We measured Ca^2+^ signals evoked by acetylcholine (1 mM, 15 s) in Chrna5+ (**Video S1**) and Syt6+ neurons (**Video S2**). To pharmacologically interrogate these populations, we measured changes in the Ca^2+^ signal and proportions of acetylcholine-responsive neurons (**Fig 2A, B**). Application of the competitive nicotinic antagonist DHBE (10 μM, 10 min) left residual acetylcholine-evoked Ca^2+^ signals in Chrna5+ neurons (35 ± 3 % of baseline, n = 71 cells, 6 mice; **Fig 2C i**) that were greater to those in Syt6+ neurons (21 ± 3% of baseline, n = 112 cells, 7 mice; Mann Whitney U = 2400, *P* < 10^-4^). The cumulative distribution of responses remaining after DHBE was significantly right-shifted in Chrna5+ neurons compared to Syt6+ neurons (**Fig 2C ii**, Kolmogorov Smirnov D = 0.37, *P* < 10^-4^). A majority of Chrna5+ neurons (~83%) still showed acetylcholine-evoked responses after DHBE, whereas fewer Syt6+ neurons (~50%) retained their responses, with a complete elimination of acetylcholine-evoked responses in the rest (**Fig 2C iii**, Fisher’s exact test: *P* < 10^-4^). Yet, the addition of muscarinic antagonist atropine did not attenuate the striking differences between Chrna5+ and Syt6+ neurons, raising the possibility of an underlying nicotinic mechanism (**Fig 2D**). In the presence of DHBE + atropine, a subset of Chrna5+ neurons still showed substantial acetylcholine-evoked Ca^2+^ signals (6 ± 1 % of baseline; **Fig 2D i**), whereas almost all Syt6+ neurons’ responses were completely blocked (0.2 ± 0.1 % of baseline; Mann Whitney U = 2083, *P* < 10^-4^) as seen from the cumulative distribution (**Fig 2D ii**, Kolmogorov Smirnov D = 0.36, *P* < 10^-4^). The proportion of Chrna5+ and Syt6+ neurons showing acetylcholine-evoked responses after DHBE+ Atropine was significantly different (41% Chrna5+ vs 6% Syt6+, Fisher’s exact test: *P* < 10^-4^, **Fig 2D iii**). Chrna5+ neurons resilient to DHBE+ Atropine within a slice appear to be localized to layer 6b (**Fig S4**). Since DHBE is a competitive antagonist, it can be outcompeted by exogenous acetylcholine at high affinity nicotinic receptors. Therefore, we hypothesized that the resilient exogenous acetylcholine-evoked responses would require non-competitive antagonist.

**Figure 2.**
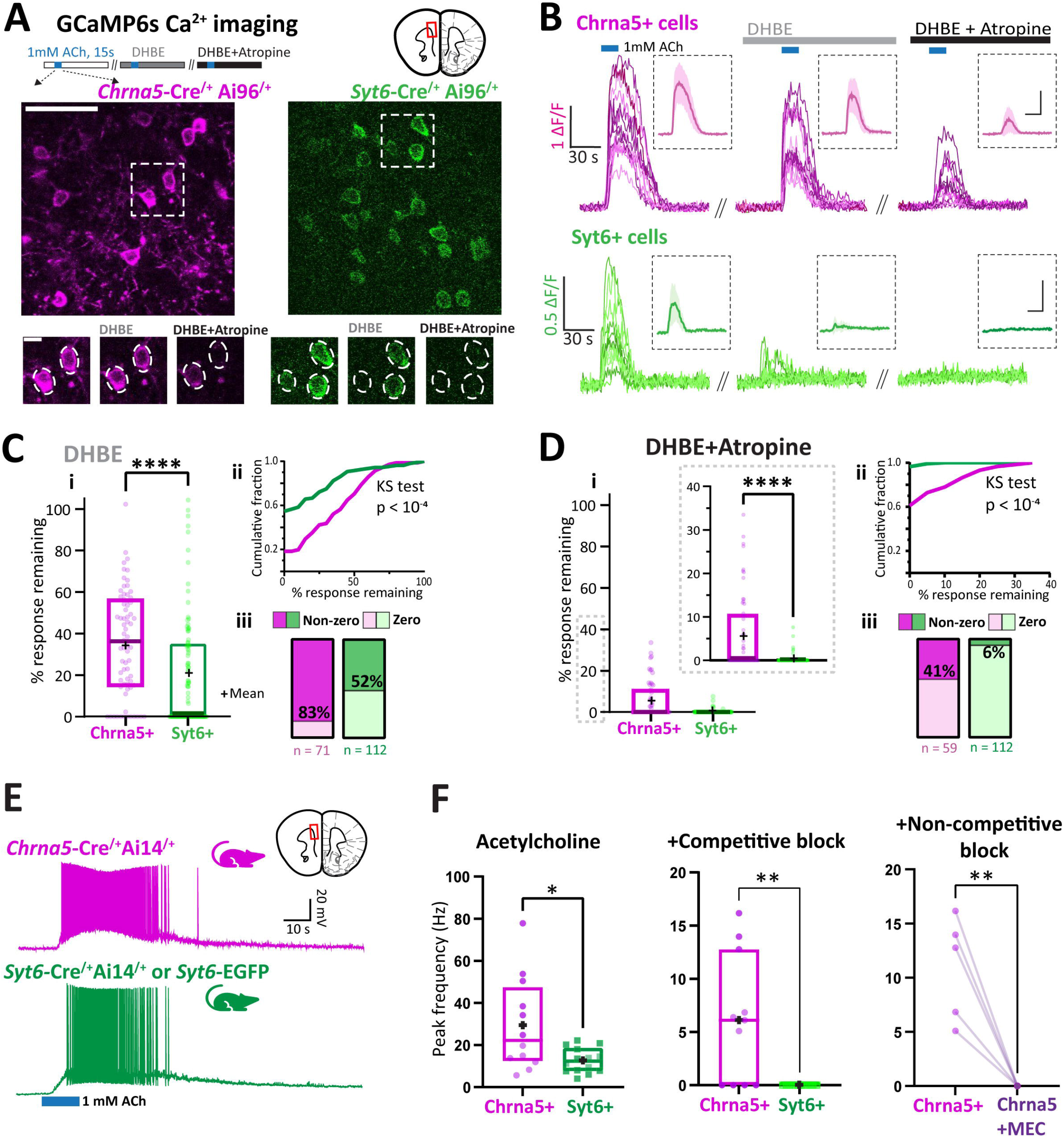
Calcium imaging in Chrna5+ and Syt6+ populations reveals a distinct subset of Chrna5+ neurons with resilient nicotinic responses. **A,** (*Top*) Two photon Ca^2+^ imaging in prefrontal brain slices from *Chrna5*-Cre^/+^Ai96^/+^ (*left*) and *Syt6*-Cre^/+^Ai96^/+^ mice (*right*) showing acetylcholine-evoked GCaMP6s responses in Chrna5+ and Syt6+ neurons respectively (scale 50 μm). (*Bottom*) Acetylcholine-evoked GCaMP6s signals were sequentially recorded after application of competitive nicotinic antagonist DHBE and addition of muscarinic antagonist atropine (scale 10 μm). **B**, Normalized fluorescence signal (ΔF by F) evoked by acetylcholine in individual Chrna5+ and Syt6+ neurons in a brain slice at baseline (*left*), after DHBE (*middle*), and after DHBE + Atropine (*right*). (Inset, average response and standard deviation. Scale: same as main figure). **C-D i,** Boxplot shows the percentage of response remaining after the application of (C) DHBE and (D) DHBE + Atropine (Inset shows the same boxplot with a restricted y-axis, ‘+’ symbols denote respective means). Responses were quantified by the area under the ΔF/F curve (n = 71 neurons, 6 mice for Chrna5+, and 112 neurons, 7 mice for Syt6+, \*\*\*\**P* <10^-4^, Mann-Whitney test). **C-D ii,** Cumulative frequency distribution of the percentage response remaining after (C) DHBE and (D) DHBE + Atropine (*P* < 10^-4^, Kolmogorov-Smirnov test). **C-D iii,** Proportion of cells showing zero and non-zero responses after (C) DHBE and (D) DHBE + Atropine (*P* < 10^-4^ for both C & D, Fisher’s exact test). **E**, Current clamp responses evoked by 1mM acetylcholine (15s) in fluorescently labeled Chrna5+ and Syt6+ layer 6 neurons patched in mPFC slices from mice *Chrna5-Cre^/+^* Ai14^/+^ and *Syt6*-Cre^/+^Ai14^/+^ or *Syt6*-EGFP mice respectively. **F**, Peak spike frequency in Chrna5+ and Syt6+ neurons evoked by acetylcholine (*left*), and in the presence of competitive nicotinic antagonist DHBE and atropine (*middle; **P* < 0.01,\**P* < 0.05, unpaired t-test). Residual response remaining after DHBE + Atropine in Chrna5+ neurons is blocked by non-competitive nicotinic antagonist mecamylamine (*right, **P* < 0.01, paired t-test). (See **Fig S4** for localization of Chrna5+ neurons in PFC with resilient nicotinic responses in layer 6b.)

To address the hypothesis of higher-affinity nicotinic binding and its consequences for spiking in these neurons, we switched to whole-cell recording in individual Chrna5+ and Syt6+ neurons. We recorded current clamp responses to acetylcholine (1 mM, 15 s) in labeled Chrna5+ and Syt6+ neurons from *Chrna5*-Cre^/+^Ai14^/+^ and *Syt6*-Cre^/+^Ai14^/+^ or *Syt6*-EGFP mice respectively (**Fig 2E**). Chrna5+ neurons showed stronger acetylcholine-evoked firing: attaining significantly higher peak firing frequency (29 ± 6 Hz, n=12 cells, 6 mice; t_(24)_ = 2.74, *P* = 0.01, unpaired t-test; Cohen’s d: 1.08) compared to Syt6+ neurons (13 ± 2 Hz, n = 14 cells, 5 mice; **Fig 2E**). The intrinsic electrophysiological properties of Chrna5+ and Syt6+ neurons did not show statistically significant differences (**Table S2**). We next examined the sensitivity of acetylcholine-evoked firing to competitive nicotinic receptor block by DHBE in the presence of atropine. Acetylcholine-evoked firing was completely eliminated in all Syt6+ neurons (**Fig 2F**) whereas a large subset of Chrna5+ neurons (7/11) retained their ability to respond to acetylcholine (Average peak firing rate in Chrna5+ neurons: 6 ± 2 Hz; t_(19)_ = 3.22, *P* = 0.004, unpaired t-test) demonstrating similar resilience to competitive nicotinic receptor block as observed with Ca^2+^ imaging. We used the non-competitive nicotinic receptor blocker mecamylamine^46,47^ to test our hypothesis that nicotinic receptors in this Chrna5+ subset were higher affinity and therefore allowed exogenous acetylcholine to outcompete DHBE. The addition of 5 μM mecamylamine was sufficient to eliminate acetylcholine-evoked firing in all the Chrna5+ neurons that were resilient to competitive nicotinic block (t_(4)_ = 5.14, *P* = 0.007, paired t-test; **Fig 2F**). Together, our Ca^2+^ imaging and electrophysiology experiments revealed the existence of a distinct subset of Chrna5+ neurons dissimilar to Syt6+ neurons, with high affinity nicotinic responses resilient to competitive nicotinic antagonism.

### Single-cell transcriptomics identifies Chrna5+ subplate neurons with Lynx genes

To determine the molecular identity of distinct Chrna5+ neurons with enhanced cholinergic responses, we pursued single-cell RNAseq. We extracted gene expression data of L5-6 glutamatergic neurons (n = 2422 cells) in the mouse anterior cingulate cortex from the Allen Institute single cell RNAseq databases^48–50^. We classified these deep-layer pyramidal neurons into 3 groups: those expressing only *Chrna5* (Chrna5+, n= 243), both *Chrna5* and *Syt6* (Chrna5+ Syt6+, n = 834), or only *Syt6* (Syt6+, n = 564) **(Fig 3A)**. 781 cells showed no expression of *Chrna5* or *Syt6* and consisted primarily of L6 Intratelencephalic cells which have been previously shown to have purely muscarinic M2/M4 mediated hyperpolarizing cholinergic responses^16,51^. We focused on the Chrna5+, Syt6+ and Chrna5+Syt6+ groups to examine their transcriptomic differences. Prior work has shown that deep cortical neurons can be separated into L6b, L5, and L6CT subclasses by the expression of characteristic markers and hierarchical clustering^50^. Single-cell analysis revealed that the Chrna5+ group primarily included L6b (44%), L5 near-projecting (L5NP, 19%), and L6CT neurons (30%), whereas the Chrna5+Syt6+ and Syt6+ groups were predominantly composed of L6CT neurons (>90%) **(Fig 3B)**. We further examined the expression of marker genes in these respective groups to validate our cell-classification. Chrna5+ neurons showed distinctive expression of several marker genes, *Ctgf, Cplx3, Kcnab1*, and *Lpar1* **(Fig 3B,C)** associated with subplate neurons^35,52^. Subplate neurons are early-born and vital for brain development, leaving L6b neurons as descendants in adulthood^36,37,53^. Notably, the highest fold enrichment among all differentially expressed genes in Chrna5+ neurons was found for subplate markers *Ctgf* (Fold change, 5.69) and *Cplx3* (3.81) (**Table 1)**. All differentially expressed genes are listed in Table S4. Overall, Chrna5+ neurons including both L5NP and L6b subpopulations highly express subplate marker genes. In contrast, *Syt6-* expressing Chrna5+Syt6+ and Syt6+ groups are only enriched in the corticothalamic markers *Foxp2* and *Syt6*, consistent with their corticothalamic subtype. These results support our imaging, electrophysiological, and pharmacological results suggesting the exclusive Chrna5+ population is a distinct cell type from typical L6CT *Syt6*-expressing neurons.

**Figure 3.**
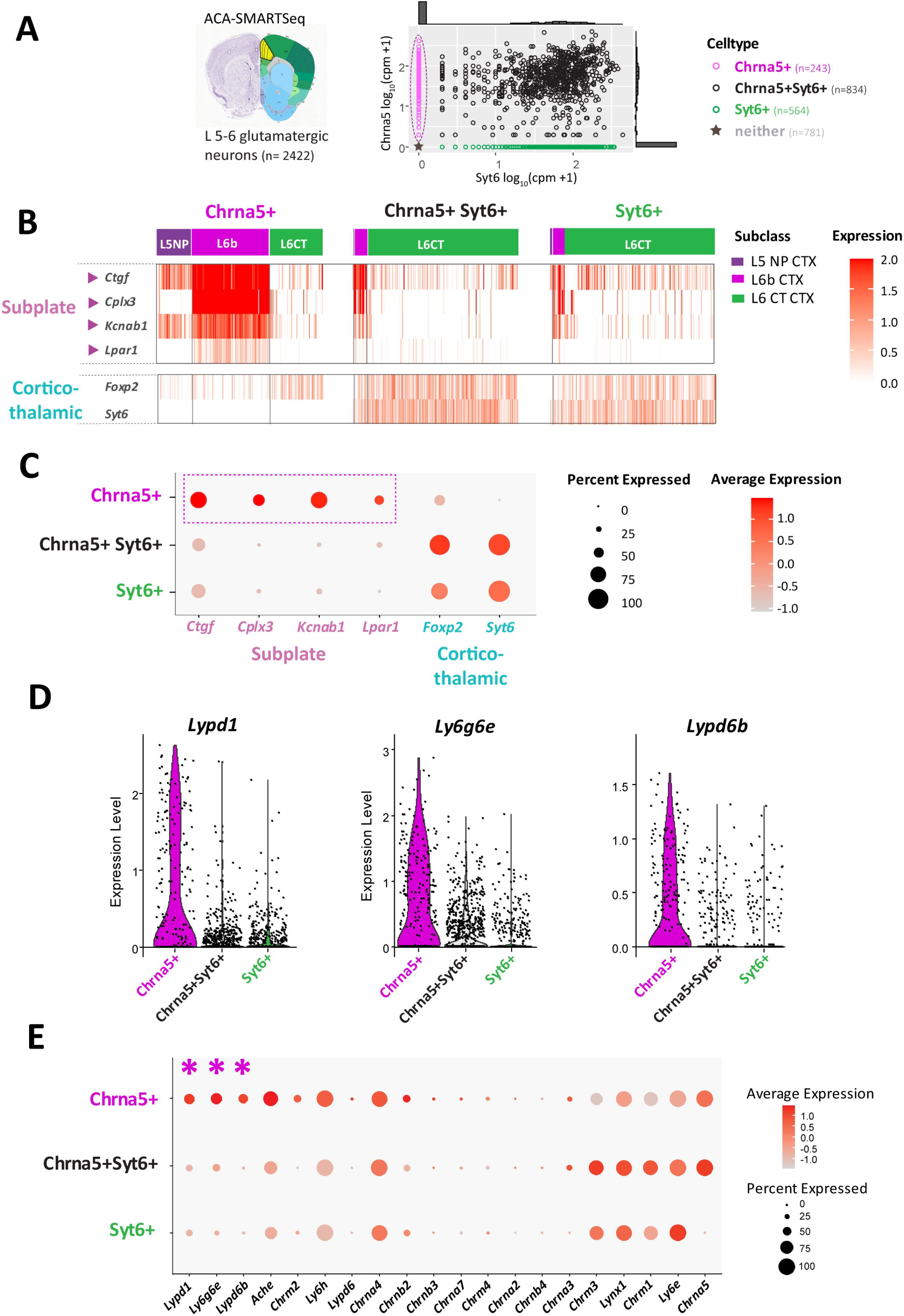
Single cell transcriptomic analysis reveals Chrna5+ subset to span subplate neuron populations with differential expression of lynx prototoxin genes. **A,** Single cell-RNAseq data for 2422 L5-6 glutamatergic neurons in the anterior cingulate cortex (ACA, shown in schematic on the left) was obtained from publicly available datasets (Allen Institute, SMARTSeq ACA and MOP (2018)). (*Right*) Scatter plot showing *Chrna5* vs *Syt6* expression in log_10_ (Copies per million +1) for each neuron, with the frequency distribution shown on the corresponding axes. Neurons were classified into Chrna5+, Chrna5+Syt6+ and Syt6+ groups based on their expression of *Chrna5* and *Syt6* genes. Cells which expressed neither gene were excluded from subsequent analyses. **B**, The major neuronal subclasses within Chrna5+, Chrna5+Syt6+ and Syt6+ groups is indicated by the colorbar on top. NP-Near projecting, CT-Corticothalamic. Heatmap shows expression of subplate and corticothalamic marker genes in each cell in all 3 groups. **C**, Dotplot shows summary of subplate and corticothalamic marker expression in Chrna5+, Chrna5+Syt6+ and Syt6+ groups. Dot size indicates the percentage of cells within each group expressing that gene, color of the dot indicates average expression level relative to other groups. Chrna5+ neurons highly express multiple subplate marker genes, but not corticothalamic markers. **D**, Violin plots show expression of Lynx prototoxin genes *Ly6g6e, Lypd1* (Lynx2) and *Lypd6b* which show highest fold-change between Chrna5+ and Chrna5+Syt6+ neurons. **E**, Dotplot shows expression of major genes known to modulate cholinergic function, including nicotinic, muscarinic subunits, acetylcholinesterase, and lynx prototoxins in Chrna5+, Chrna5+Syt6+ and Syt6+ neurons. Genes are ordered by decreasing fold change in expression. Dot size indicates the percentage of cells within each group expressing that gene, color of the dot indicates average expression level relative to other groups. Fold change of all the genes shown in this dotplot are listed in Table S3. (See **Fig S5** for transcriptomic analysis of Chrna5+ neurons in other cortical areas showing conserved subplate markers and lynx prototoxin gene expression across cortical regions)

**Table 1.**
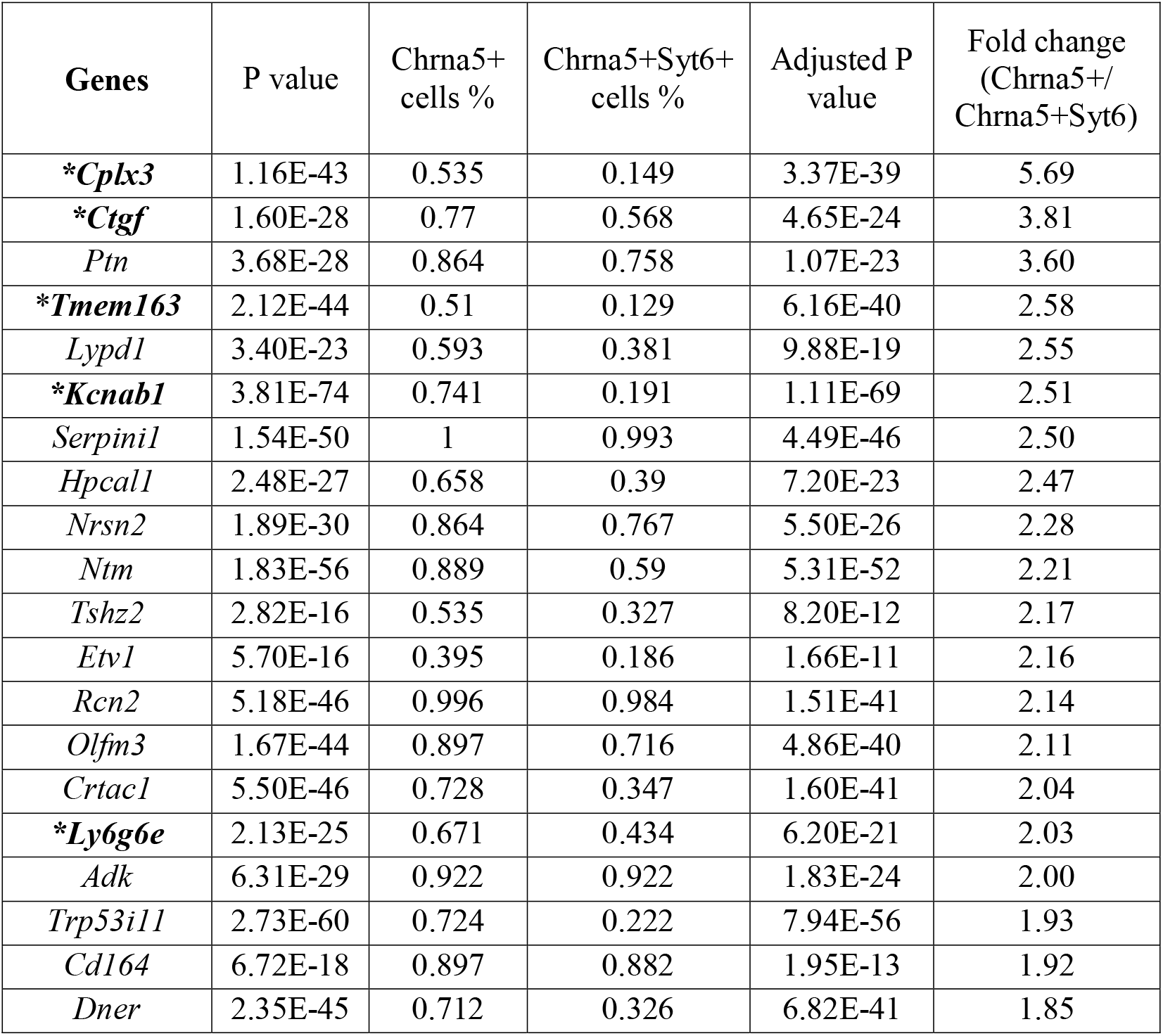
Top 20 differentially expressed genes between Chrna5+ and Chrna5+Syt6+ neurons. Top 20 differentially expressed genes determined by the FindMarkers function on Seurat ordered by decreasing fold change. Several known subplate neuron markers (highlighted by *) are the highest enriched genes in Chrna5+ neurons. All differentially expressed genes are listed in supplementary table S4.

| Genes | P value | Chrna5+<br>cells % | Chrna5+Syt6+<br>cells % | Adjusted P<br>value | Fold change<br>(Chrna5+/<br>Chrna5+Syt6) |
| --- | --- | --- | --- | --- | --- |
| **\*Cplx3** | 1.16E-43 | 0.535 | 0.149 | 3.37E-39 | 5.69 |
| **\*Ctgf** | 1.60E-28 | 0.77 | 0.568 | 4.65E-24 | 3.81 |
| *Ptn* | 3.68E-28 | 0.864 | 0.758 | 1.07E-23 | 3.60 |
| **\*Tmem163** | 2.12E-44 | 0.51 | 0.129 | 6.16E-40 | 2.58 |
| *Lypd1* | 3.40E-23 | 0.593 | 0.381 | 9.88E-19 | 2.55 |
| **\*Kcnab1** | 3.81E-74 | 0.741 | 0.191 | 1.11E-69 | 2.51 |
| *Serpini1* | 1.54E-50 | 1 | 0.993 | 4.49E-46 | 2.50 |
| *Hpcal1* | 2.48E-27 | 0.658 | 0.39 | 7.20E-23 | 2.47 |
| *Nrsn2* | 1.89E-30 | 0.864 | 0.767 | 5.50E-26 | 2.28 |
| *Ntm* | 1.83E-56 | 0.889 | 0.59 | 5.31E-52 | 2.21 |
| *Tshz2* | 2.82E-16 | 0.535 | 0.327 | 8.20E-12 | 2.17 |
| *Etv1* | 5.70E-16 | 0.395 | 0.186 | 1.66E-11 | 2.16 |
| *Rcn2* | 5.18E-46 | 0.996 | 0.984 | 1.51E-41 | 2.14 |
| *Olfm3* | 1.67E-44 | 0.897 | 0.716 | 4.86E-40 | 2.11 |
| *Crtac1* | 5.50E-46 | 0.728 | 0.347 | 1.60E-41 | 2.04 |
| **\*Ly6g6e** | 2.13E-25 | 0.671 | 0.434 | 6.20E-21 | 2.03 |
| *Adk* | 6.31E-29 | 0.922 | 0.922 | 1.83E-24 | 2.00 |
| *Trp53i11* | 2.73E-60 | 0.724 | 0.222 | 7.94E-56 | 1.93 |
| *Cd164* | 6.72E-18 | 0.897 | 0.882 | 1.95E-13 | 1.92 |
| *Dner* | 2.35E-45 | 0.712 | 0.326 | 6.82E-41 | 1.85 |

To identify molecular changes predictive of Chrna5+ nicotinic ‘super-responders’, we examined differential expression of genes that exert effects on postsynaptic cholinergic responses (**Fig 3D-E**). We selected cholinergic receptor genes (nicotinic *Chrna5-2, Chrna7, Chrnb2-4*, and muscarinic subunits *Chrm1-4*) acetylcholinesterase (*Ache*), and members of the family of genes that encode lynx proteins (*Ly6e, Ly6h, Ly6g6e, Lynx1, Lypd1, Lypd6, Lypd6b*) known to allosterically modulate nicotinic receptor responses^38^. We found substantial and highly significant changes in the expression of three lynx prototoxins *Lypd1, Ly6g6e* and *Lypd6b* **(Fig 3D)**. While both Chrna5+ and Chrna5+Syt6+ populations express *Chrna5*, there was slightly higher expression of *Chrna5* (25% increase) as well as lower expression of the inhibitory muscarinic receptor *Chrm2* (20% decrease) in Chrna5+Syt6+ neurons. There were no significant differences in other nicotinic and muscarinic subunit expression between the two groups. Acetylcholinesterase, the enzyme that breaks down acetylcholine was also highly expressed (50% increase) in Chrna5+ neurons, which may benefit their nicotinic responses by protecting receptors from overactivation and desensitization. Of note, *Chrnb2* expression does seem to be in fewer cells than expected (Fig 3E), which can be attributed to sparse detection of *Chrnb2* levels by scRNAseq approaches^54,56^. The fold-change of the genes in **Fig 3E** between Chrna5+ and Chrna5+Syt6+ neurons is shown in **Table S3**. Notably, the top three genes with highest fold change in Chrna5+ neurons were the GPI-anchored lynx prototoxins: Lynx2 encoded by *Lypd1* (Fold change: 2.55), *Ly6g6e* (2.03), and *Lypd6b* (1.51). The distinct pattern of expression of specific lynx proteins in Chrna5+ neurons suggests unexpectedly complex endogenous control of nicotinic responses in these prefrontal subplate neurons.

Transcriptomic examination of *Chrna5*-expressing neurons in other cortical regions including primary motor (MOp), somatosensory (SSp), and visual cortices (VISp) also confirmed that Chrna5+ neurons consist of L6b and L5/6NP neurons expressing subplate marker genes, in addition to the lynx modulatory proteins as described above in the PFC (**Fig S5**). Therefore, *Chrna5-expression* identifies a conserved population of deep-layer 6 neurons with subplate identity across multiple cortical regions that display specialized expression of lynx proteins to modulate nicotinic receptors.

### Perturbing native prefrontal cortical lynx-modulation of optogenetic nicotinic responses

To examine whether the molecular machinery of deep layer prefrontal neurons endows them with greater dynamic range in responding to acetylcholine, we sought to experimentally perturb endogenous lynx modulation. Members of the lynx-family are GPI-anchored (**Fig 4A**), and work in cell expression systems^39^ suggests these anchors can be cleaved via activation of phospholipase C (PLC). These experiments are important because the potential impact of GPI cleavage on nicotinic responses in a native system is not well understood. We hypothesized that perturbing lynx-mediated control could affect endogenous nicotinic properties in a complex manner (**Fig 4A**) since both positive (eg. Ly6g6e) and negative modulatory lynx proteins (eg. Lynx1) are expressed. To cleave GPI-anchored proteins, we used the PLC activator compound m-3M3FBS^55,57–59^. Nicotinic responses of deep layer pyramidal neurons from ChAT-ChR2 mice to optogenetic acetylcholine release were recorded in the continuous presence of atropine before and after treatment with m-3M3FBS (25μM, 5 min; **Fig 4B**). The rising slope of the nicotinic responses showed a significant increase after m-3M3FBS treatment (23 ± 17%; Paired Cohen’s d = 0.83; *P* = 0.008, Wilcoxon matched-pairs test), compared to the baseline change observed in the same cells prior to PLC activation (−6 ± 4%; 8 cells, 6 mice; **Fig 4C, D**). This increase was not observed with the inactive ortholog o-3M3FBS that does not activate PLC (Paired Cohen’s d = 0.09, *P* = 0.625, Wilcoxon matched-pairs test, data not shown). The area under the nicotinic response known as charge transfer also showed a statistically significant increase following PLC activation (22 ± 7%; Cohen’s d = 1.68; *P* = 0.016, Wilcoxon matched-pairs test; **Fig 4D**), compared to baseline change (−9 ± 6%). Thus, PLC activation causes a specific increase in nicotinic receptor responses, presumably due to cleavage of inhibitory GPI-anchored prototoxins such as Lynx1.

**Figure 4.**
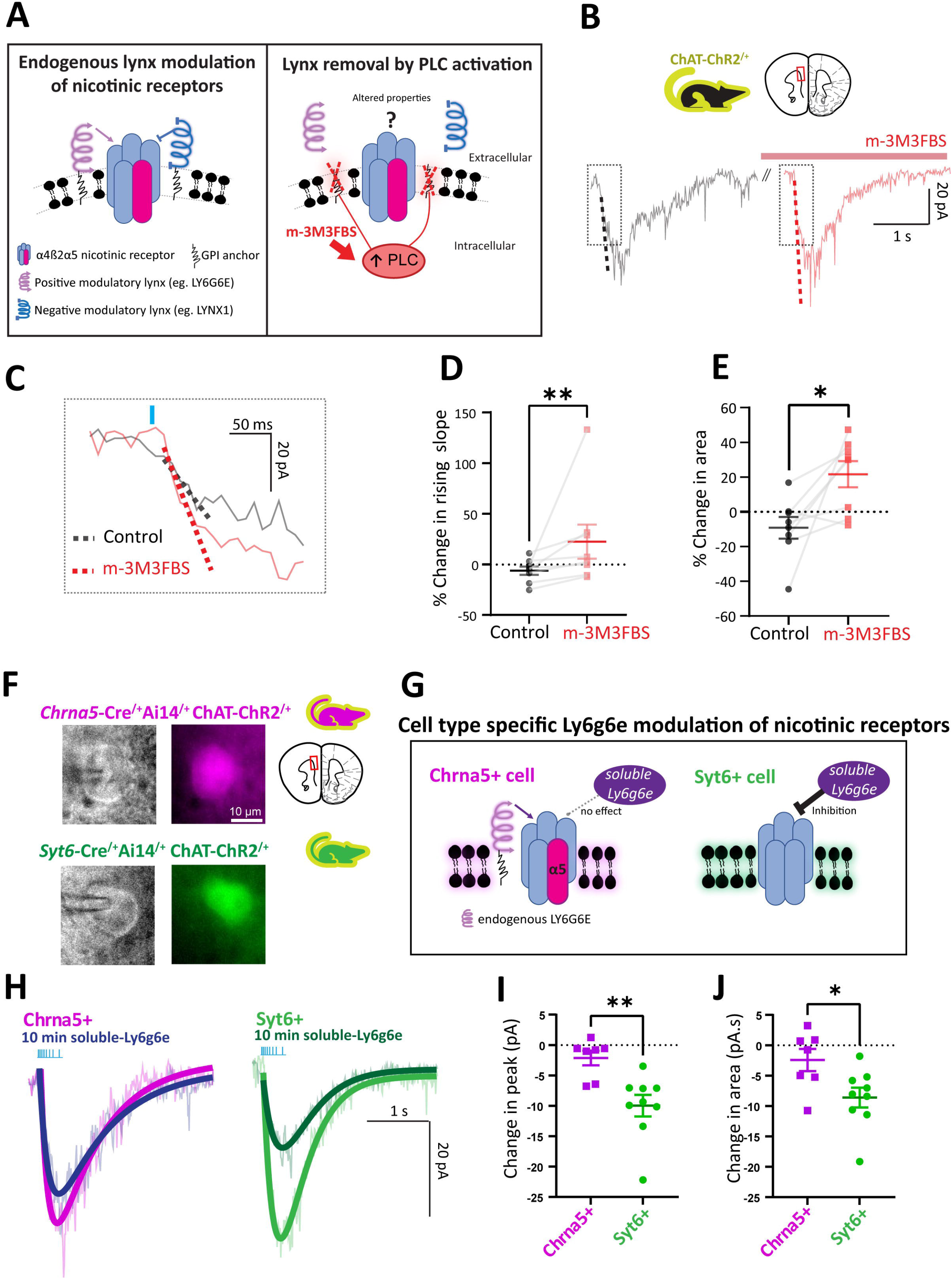
Regulation of optogenetic nicotinic responses by endogenous GPI-anchored lynx proteins and cell type specific effects of recombinant Ly6g6e. **A,** Schematic of nicotinic receptor environment showing endogenous GPI-anchored lynx proteins exerting positive and negative modulation of nicotinic receptors. The compound m-3M3FBS activates PLC, cleaving the GPI anchor and perturbing lynx-mediated modulation of nicotinic responses. **B,** Optogenetic nicotinic responses in prefrontal deep-layer pyramidal neurons from ChAT-ChR2 mice before and after treatment with m-3M3FBS (5 min). **C,** PLC activation significantly increased the rising slope of optogenetic nicotinic responses. Percent change in **D,** Rising slope and **E,** Area of nicotinic response in control and after PLC activation. (\**P* < 0.05, \*\**P* < 0.01, Wilcoxon matched-pairs test). **F**, IRDIC (*left*) and widefield fluorescence (*right*) images of tdTomato labeled Chrna5+ (*top*) and Syt6+ (*bottom*) neurons during whole-cell patch clamp electrophysiology in *Chrna5*-Cre^/+^Ai14^/+^ChAT-ChR2^/+^ and *Syt6*-Cre^/+^Ai14^/+^ChAT-ChR2^/+^ mouse brain slices respectively. **G,** Schematic summarising predicted and observed effects of recombinant water-soluble ly6g6e on Chrna5+ and Syt6+ neuronal nicotinic receptors. **G,** Optogenetic nicotinic responses are reduced in amplitude following 10 minute application of soluble ly6g6e in Syt6+ but not Chrna5+ neurons. Change in peak current (**I**) and area of the nicotinic response (**J**) of Chrna5+ vs Syt6+ neurons (\**P* < 0.05, ** *P* < 0.01, Unpaired t-test)

To test the transcriptomic prediction that cell-type specific differences in lynx modulation lead to different cholinergic properties, we obtained purified water-soluble recombinant Ly6g6e protein and examined its effects on optogenetic nicotinic responses in labeled Chrna5+ and Syt6+ neurons (**Fig 4F**). These experiments were conducted in *Chrna5*-Cre^/+^Ai14^/+^ChAT-ChR2^/+^ and *Syt6*-Cre^/+^Ai14^/+^ChAT-ChR2^/+^ mice. We hypothesized that the modulation of Chrna5+ neuronal nicotinic receptors by endogenous Ly6g6e would occlude the effect of exogenous soluble Ly6g6e, whereas Syt6+ neurons would be altered by exposure to the exogenous Ly6g6e (Fig 4G). Consistent with this hypothesis, we found that 10 minute application of soluble Ly6g6e did not significantly alter the amplitude of optogenetically evoked nicotinic responses in labeled Chrna5+ neurons (Change in peak = −2.1 ± 1.2 pA, t_(6)_ = 1.79, *P* = 0.12, paired t-test). However, in labeled Syt6+ neurons lacking endogenous expression of *Ly6g6e*, exogenous application of soluble Ly6g6e caused a significant decrease in the amplitude (Change in peak = −10 ± 1.8 pA, t_(8)_ = 5.60, *P* < 0.001, paired t-test; **Fig 4H-I**). The change in peak and area of the nicotinic responses caused by solube Ly6g6e was significantly different between Chrna5+ and Syt6+ neurons (change in peak: t_(14)_ = 3.43, *P* = 0.004; Change in area: t_(14)_ = 2.53, *P* = 0.024, Unpaired t test; **Fig 4I-J**). Of note, soluble and endogenous GPI-anchored prototoxins are known to have opposite effects on nicotinic receptors and the exact direction of endogenous modulation of nicotinic receptors by different lynx proteins is still debated^41,60,61^. The key outcome of this experiment is the difference in the Ly6g6e modulation of Chrna5+ and Syt6+ neurons, not the direction. We conclude that Chrna5+ neurons exert specialized molecular control over their nicotinic receptors, shaping their fate as acetylcholine super-responders.

## Discussion

Our work examines the effects of GPI-anchored lynx prototoxins on native nicotinic receptor-mediated optogenetic responses, advancing from work in heterologous expression systems. These results are a first step in showing how endogenous lynx regulation of nicotinic responses can act in a complex cell-type specific fashion leading to specialized cholinergic properties in a subset of neurons. Overall, our study reveals a previously uncharacterized population of *Chrna5-* expressing subplate neurons in the prefrontal cortex that are exquisitely sensitive to acetylcholine, with differential expression of several lynx prototoxin genes that allow flexible tuning of their high-affinity nicotinic responses (**Fig 5**).

**Figure 5.**
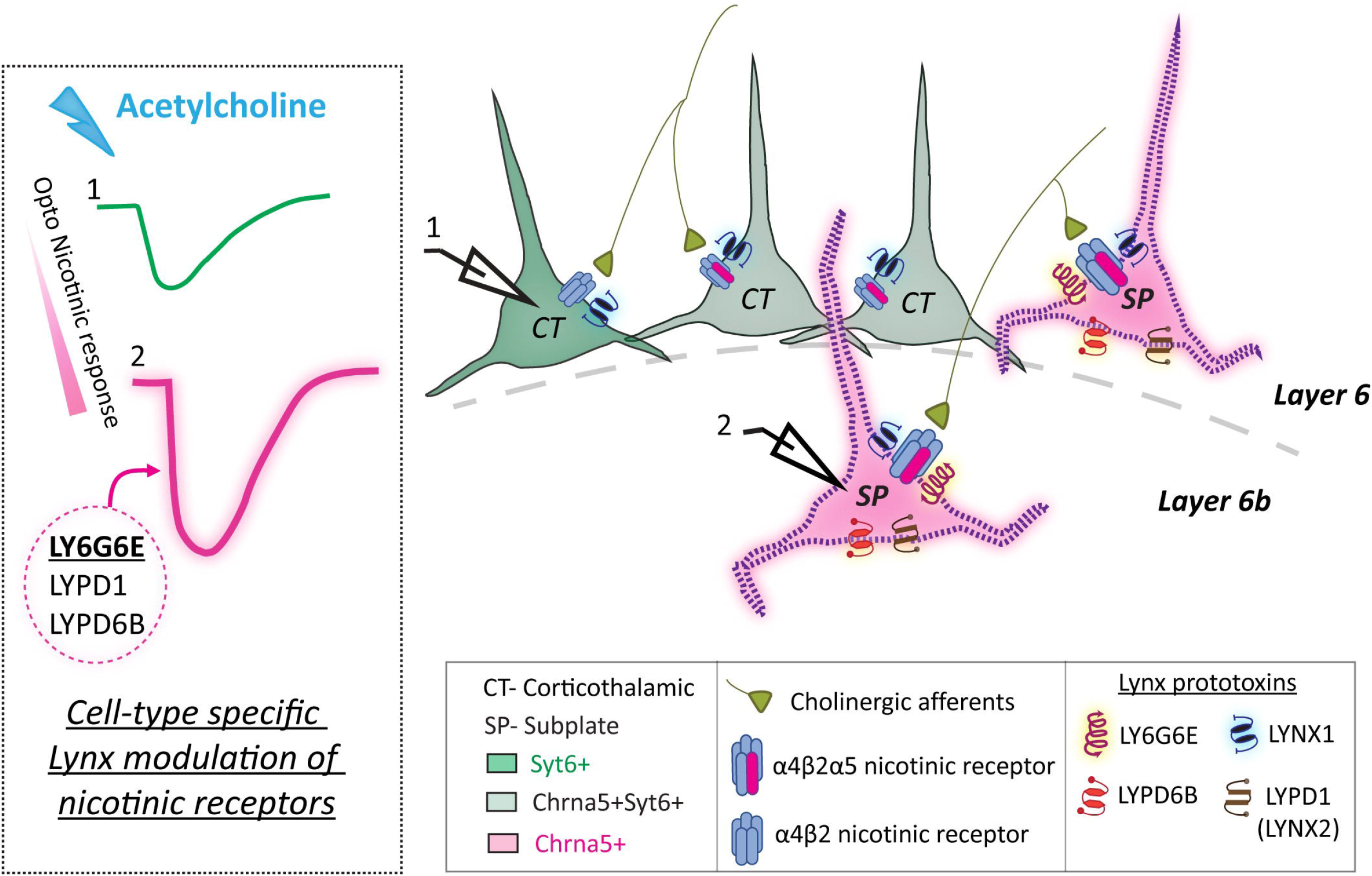
Graphical summary. Deep-layer pyramidal neurons can be divided into three groups (Chrna5+, Chrna5+Syt6+, Syt6+) by their expression of *Chrna5* and *Syt6* genes. The subset of *Chrna5-expressing* neurons without *Syt6* expression are molecularly distinct and comprise of subplate neurons, whereas *Syt6*-expressing neurons are of the corticothalamic subtype. Nicotinic receptors in these neurons are under complex regulation by endogenous lynx prototoxins. Inhibitory prototoxin gene *Lynx1* is expressed uniformly in all neurons, whereas Chrna5+ subplate neurons additionally have specific expression of *Ly6g6e, Lypd1* and *Lypd6b* prototoxin genes. These Chrna5+ subplate neurons in layer 6b show enhanced α5 subunit nicotinic receptor-mediated cholinergic responses that are differently modulated by specific lynx prototoxins.

### Specialized cholinergic properties of Chrna5-expressing neurons

While important for attention, the neurophysiological impact of the auxiliary α5 nicotinic subunit in its native neuronal environment has remained beyond the reach of previous work. The contributions of α5 to high-affinity nicotinic receptors have been extrapolated based on results of cell system experiments and work in rodents deleted for *Chrna5*^13,16,22,62,63^. Here, *Chrna5-Cre* mice allowed us to affirmatively demonstrate that neurons expressing the α5 nicotinic subunit respond faster and more strongly to endogenous acetylcholine. This cholinergic heterogeneity among layer 6 neurons prompted a larger scale comparison between Chrna5+ neurons and a well-defined layer 6 population labeled by *Syt6*^29^. These experiments revealed a subset of Chrna5+ acetylcholine ‘super-responders’ with high affinity nicotinic responses that were not found in Syt6+ neurons.

### Heterogeneity of cell types expressing Chrna5

Previously, the deep-layer cell types expressing *Chrna5* were uncharacterized, and generally thought to include L6CT neurons^8^. Investigation of L6CT neurons have relied on *Syt6*-Cre and *Ntsr1-Cre* mouse lines that label similar sets of neurons^27–29^, with only *Syt*6-Cre mice successfully labeling this population in prefrontal cortex^31^. L6CT neurons labeled by *Syt6* or *Ntsr1* expression are excited by acetylcholine^32,64^, but the degree to which their nicotinic response relied on *Chrna5* expression was unclear. Strikingly, we reveal that acetylcholine super-responders with high affinity nicotinic receptors are from the population of Chrna5+ neurons without *Syt6-*expression. Transcriptomic analysis demonstrates that majority of these likely Chrna5+ ‘super-responders’ arise from L5 Near-Projecting and L6b neurons, populations that express multiple markers linking them to the developmental subplate. These enigmatic cells are remnants of earliest-born cortical neurons that serve as a relay for establishing connections between cortex and thalamus^65,66^. Subplate neurons receive cholinergic inputs at birth^67^, highlighting their role in developmental cholinergic modulation.

### Advantages of Chrna5 as a marker for subplate cells

In contrast to L6CT neurons, subplate neurons remain relatively uncharacterized due to the lack of transgenic mice to definitively label the diverse subtypes, and inaccessibility of the available lines for *in vivo* targeting. Transcriptomic analysis (Fig 5, Table 1) suggests that the Chrna5+ population is enriched for known subplate markers *Ctgf* (Connective tissue growth factor), *Cplx3* (Complexin 3), *Kcnab1*,and *Lpar1*^35,68,69^. Significantly, the lynx prototoxin and nicotinic receptor modulator *Ly6g6e*, which is highly expressed in Chrna5+ neurons, is also a marker of subplate neurons^70^. Our study is the first to identify enhanced cholinergic activation regulated by *Chrna5* and lynx-gene expression in subplate/L6b neurons. Subplate neurons have recently been found to strongly regulate cortical output through their intracortical connections^71,72^. Enhanced cholinergic activation in these neurons will have different consequences for prefrontal processing, challenging the popular conception that cholinergic modulation of attention occurs only through top-down control of thalamic input by L6CT neurons.

### Molecular determinants of nicotinic receptor properties in Chrna5+ neurons

Our transcriptomic analysis revealed enhanced expression of GPI-anchored lynx prototoxin genes *Ly6g6e, Lypd1*, and *Lypd6b* in Chrna5+ neurons (Fig 5). Lynx proteins are well known modulators of nicotinic receptor properties and trafficking^41,43^, but most of the insight into their actions comes from heterologous cell systems, deletion, and overexpression experiments. Their effects on nicotinic receptors in their native environment is unclear. In expression systems, Ly6g6e potentiates α4β2 nicotinic responses, slowing their desensitization^39^, predicting cholinergic responses in Chrna5+ neurons would be resistant to desensitization as has been implied by *Chrna5* deletion work^16^. In contrast, Lynx2 (*Lypd1*) is a predicted negative modulator that can increase desensitization of α4β2 nicotinic receptors^73^. Lynx2 acts intracellularly to reduce surface expression of α4β2 nicotinic receptors^39^, but may preferentially act on lower affinity (α4)_3_(β2)_2_ receptors^62,74^ and indirectly promote expression of high affinity (α4)_2_(β2)_2_α5 nicotinic receptors. The effect of Lypd6b on (α4)_2_(β2)_2_α5 nicotinic receptors is yet to be determined and may further contribute to the complex control of Chrna5+ nicotinic responses^75^. In addition, Lynx1, a well-known negative modulator of α4β2 nicotinic receptors^42,76–78^ is also expressed in Chrna5+ neurons. Consistent with such complex lynx regulation, our experiments confirmed that removing GPI-anchored lynx proteins increases nicotinic response onset and amplitude potentially due to removal of Lynx1. In contrast, exogenous application of recombinant Ly6g6e had different effects in Chrna5+ and Syt6+ neurons, consistent with cell-type specific lynx modulation in Chrna5+ neurons predicted by transcriptomics.

### Functional consequences

The effects of lynx proteins on nicotinic receptor function have so far been determined by heterologous expression systems^39^, knockout studies^73,79^, exogenous application of recombinant water-soluble lynx proteins^80,81^, and viral manipulation of expression in the brain^42,82^. We advance this field by revealing, in native tissue, complex endogenous regulation of optogenetic nicotinic responses by multiple GPI-anchored lynx proteins. Inhibitory lynx expression and high levels of acetylcholinesterase in Chrna5+ neurons suggest that their responses are restrained and our experiments likely underestimated their nicotinic receptor function. These responses could be dramatically enhanced when acetylcholinesterase and inhibitory lynx modulation is reduced through other signaling mechanisms. Such flexible tuning of nicotinic responses by lynx prototoxins in Chrna5+ neurons can provide greater dynamic range and poises them to be key players during attentional processing (Fig 5). A recent study found that preventing developmental increase in Lynx1 expression in corticocortical neurons by viral knockdown led to altered cortical connectivity and impaired attention^42^. Thus cell-type specific changes in lynx expression during development are critical for maturation of attention circuits. It is of interest to examine such changes during development in Chrna5+ neurons and how they differ from Syt6+ neurons.

### Caveats and limitations of study

Subplate marker genes and localization of subplate cells can vary across cortical regions, and they have been better studied in sensory cortical regions^83–85^. Our optophysiological, anatomical and transcriptomic data show Chrna5+ acetylcholine super-responder neurons to be found in layer 6b both in the primary somatosensory cortex and PFC (Fig S2-S3). However, an outstanding question is whether transcriptomic cell classes such as L5/6NP and L6b, which express multiple subplate marker genes, correlate with early developmental subplate origin. Anatomical evidence alone is insufficient for this purpose. For example, while the distinct transcriptomic subclasses L6b and L5/6NP show overlapping spatial distribution in deep-layers of the motor cortex^86,87^ these subclasses possess distinct epigenetic signatures whose implications are yet to be understood. It will be necessary for future work to examine whether transcriptomic /epigenetically defined cell classes (L5/6NP, L6b) expressing subplate markers also show distinct early developmental origin. Lack of labeling strategies has limited efforts to characterize these neuronal subclasses and verify their developmental subplate identity. *Chrna5-Cre* mice will be an important tool to rectify this as they consistently label these conserved subclasses across multiple cortical regions. Another caveat of our study is the perturbation of lynx prototoxins using chemical strategies. While we have demonstrated cell-type specific effects of recombinant ly6g6e, it remains to be seen whether cell-type specific expression of ly6g6e or other lynx prototoxins is necessary for nicotinic modulation. Future work is needed to genetically manipulate lynx prototoxins in specified cell populations.

### Summary of advances

Our study reveals a distinct group of ‘acetylcholine super-responder’ neurons in the prefrontal cortex identified by *Chrna5-expression* that constitute subplate neurons vital for cortical development. We identify that their high affinity α5 subunit-containing nicotinic receptors are under complex regulation by several lynx prototoxins and acetylcholinesterase. *Chrna5-Cre* mice are a valuable tool for future studies examining the in vivo role of these specialized neurons.

## Acknowledgements

We acknowledge the generous support of the Canadian Institutes of Health Research (CIHR MOP 89825, EKL; CIHR PJT-153101, EKL), the Canada Research Chair in Developmental Cortical Physiology (EKL), Ontario Graduate Scholarships (SV). We thank Ms. Janice McNabb and Mr. Ha-Seul Jeoung for expert technical assistance. Earlier presentations of this work received valuable feedback from Dr. Junchul Kim, Dr. Beverly Orser, and Dr. Steve Prescott of the University of Toronto.

## STAR Methods

### Resource availability

#### Lead contact

Further information and requests for resources and reagents should be directed to Dr. Evelyn Lambe or Dr. Eric Turner.

#### Materials availability

This study did not generate new unique reagents.

#### Data and code availability

This paper analyzes existing, publicly available single cell RNAseq data. The links for the Allen Institute scRNAseq database are provided in the key resources table. The paper does not report original code. Any additional information required to reanalyze the data reported in this paper is available from the lead contact upon request.

### Experimental model and subject details

*Syt6*-Cre^/+^GCaMP6s^/+^ and *Chrna5*-Cre^/+^GCaMP6s^/+^ mice used for Ca^2+^ imaging were obtained by crossing *Chrna5-Cre* (Gift from Dr. Eric Turner) and *Syt6*-Cre mice (*Syt6*-Cre KI148, RRID:MMRRC 037416-UCD,^90^) respectively, with Ai96 mice (JAX: 024106). For electrophysiological recordings of labeled Chrna5+ and Syt6+ neurons, we used *Chrna5-* Cre^/+^Ai14^/+^, and *Syt6*-Cre^/+^Ai14^/+^ mice respectively. *Syt6*-EGFP^/+^ mice were additionally used for few experiments (*Syt6*-EGFP EL71, RRID:MMRRC 010557-UCD,^91^).

Triple transgenic mice labeling both *Chrna5* and *Syt6-*expressing neurons with EGFP in Syt6+ neurons and tdTomato in Chrna5+ neurons were used to examine the overlap between the two cell types. *Syt6*-EGFP and Ai14 mice^92^ were bred together and the offspring were crossed with *Chrna5-Cre* mice to generate *Chrna5*-Cre^/+^Ai14^/+^*Syt6*-EGFP^/+^ mice used for these experiments. A set of experiments measuring optogenetic cholinergic responses was also performed in ChAT-ChR2 (ChAT^/+^) mice (JAX: 014546). To examine optogenetic cholinergic responses in labeled Chrna5 and Syt6 cell populations, the respective Cre lines were crossed with ChAT^/+^Ai14^/+^ mice to generate *Chrna5*-Cre^/+^Ai14^/+^ChAT^/+^ and *Syt6*-Cre^/+^Ai14^/+^ChAT^/+^ mice.

All animals were bred on a C57BL/6 background, except *Syt6*-EGFP which were Black Swiss. Adult male and female animals age >P60 were used in the study. Mice were separated based on sex after weaning at P21 and group-housed (2-4 mice per cage). Animals had ad libitum access to food and water and were on a 12-h light/dark cycle with lights on at 7 AM. Guidelines of the Canadian Council on Animal Care were followed, and all experimental procedures were approved by the Faculty of Medicine Animal Care Committee at the University of Toronto. 42 mice were used for the entire study, with similar numbers of males and females.

### Method details

#### Brain slicing and electrophysiology

Slicing and electrophysiology followed procedures described previously^16^. An intraperitoneal injection of chloral hydrate (400 mg/kg) was given to anesthetize mice prior to decapitation. The brain was rapidly extracted in ice cold sucrose ACSF (254 mM sucrose, 10 mM D-glucose, 26 mM NaHCO_3_, 2 mM CaCl_2_, 2 mM MgSO_4_, 3 mM KCl and 1.25 mM NaH_2_PO_4_). 400 μm thick coronal slices of prefrontal cortex (Bregma 2.2 − 1.1) were obtained on a Dosaka linear slicer (SciMedia, Costa Mesa, CA, USA). Slices were left to recover for at least 2 hours in oxygenated (95% O2, 5% CO_2_) ACSF (128 mM NaCl, 10 mM D-glucose, 26 Mm NaHCO_3_, 2 mM CaCl_2_, 2 mM MgSO_4_, 3 Mm KCl, and 1.25 mM NaH_2_PO_4)_ at 30°C before being used for electrophysiology or two-photon Ca^2+^ imaging. Brain slices were transferred to the stage of a BX51WI microscope (Olympus, Tokyo, Japan) and perfused with oxygenated ACSF at 30°C. Recording electrodes (2 - 4 MΩ) containing patch solution (120 mM potassium gluconate, 5 mM KCl, 10 mM HEPES, 2 mM MgCl_2_, 4 mM K_2_-ATP, 0.4 mM Na_2_-GTP and 10 mM sodium phosphocreatine, pH adjusted to 7.3 using KOH) were used to patch pyramidal neurons in layer 6 - 6b based on morphology and proximity to white matter. Only regular spiking neurons were included. Multiclamp 700B amplifier at 20 kHz with Digidata 1440A and pClamp 10.7 software (Molecular devices) were used for data acquisition. All recordings were compensated for liquid junction potential (14 mV). Voltage-clamp responses were examined at −75 mV and in current-clamp at rest or starting from −70 mV.

#### Optogenetics

5 ms pulses of blue light (473 nm) were delivered through the 60X objective lens with an LED (Thorlabs, 2 mW) to excite channelrhodopsin containing cholinergic fibers. Pattern of stimulation was as in a previous study, with 8 pulses of light delivered in a frequency accommodating manner^16^.

#### Pharmacology

Acetylcholine (1mM, Sigma) was used to exogenously stimulate cholinergic receptors. Atropine (200 nM, Sigma) and Dihydro-β-erythroidine (DHBE, 10 μM, Tocris) were used to competitively block muscarinic receptors and β2 subunit-containing nicotinic receptors respectively. Mecamylamine (5 μM, Tocris) was used to further non-competitively block nicotinic receptors. Phospholipase C activator m-3M3FBS (25 μM, Tocris) was used to cleave GPI-anchored Lynx prototoxins and the inactive ortholog o-3M3FBS (25 μM, Tocris) was used as a control^55^. Cyclodextrin (1 mM, Tocris) was included in a small subset of experiments to improve solubility of 3M3FBS compounds, but no further improvement in efficacy was observed. Water soluble recombinant Ly6g6e (0.5 mg/ml) was obtained by custom purification (Creative Biomart) and used for exogenous application at 1: 1000 and 3:1000 dilution. Effects on nicotinic receptors were not distinguishable between the two different protein concentrations. Only freshly thawed protein aliquots were used for experiments.

#### Two-photon imaging

Two-photon imaging of GCaMP6s Ca^2+^ signals in L6 neurons was performed using a 60× water-immersion objective with 0.90 numerical aperture using an Olympus Fluoview FV1000 microscope and a Titanium-Sapphire laser sapphire laser (Newport) at 930nm. Images were sampled at 512 x 512 pixels (2.4 pixels/μm) at a frame rate of 0.9 Hz. Following a 2-minute washout period for this initial application, GCaMP6s Ca^2+^ signals were measured in response to acetylcholine (1 mM, 15 s). The cellular responses to acetylcholine were measured at baseline, then after application of competitive nicotinic receptor antagonist DHBE (10 μM, 10 min), and again after the addition of muscarinic antagonist atropine (200 nM, 10 min).

Dual color two-photon imaging (910 nm excitation, using 570 nm dichroic mirror with green (540-595 nm) and red (570-620 nm) filters) was performed in brain slices of triple transgenic *Chrna5*-Cre^/+^Ai14^/+^*Syt6*-EGFP^/+^ mice to examine overlap in fluorescent reporter expression between Chrna5+ and Syt6+ neurons. Z-stacks of 30 frames acquired in 1-μm steps were taken in layer 6 of mPFC slices and the maximum projection used to count cells with the cell counter feature in Fiji. A set of mPFC brain slices from *Chrna5*-Cre^/+^Ai14^/+^*Syt6*-EGFP^/+^ mice were also fixed and mounted for confocal imaging with LSM880 (Leica) microscope.

#### Single cell RNAseq analysis

Single cell RNAseq data for Anterior Cingulate Cortex (ACA) of adult mice was taken from the AC A and MOP Smart-Seq (2018) database, with cell-type annotations from Whole cortex & Hippocampus Smart-Seq (2019) database from the Allen Institute for Brain Science at https://portal.brain-map.org/atlases-and-data/rnaseq^48,50^. Single cell analysis was performed using the R package Seurat (v 4.04)^93–95^. Layer 5 and 6 glutamatergic neurons were selected and sorted into three cell classes based on their expression of *Chrna5* and *Syt6* genes: those expressing only *Chrna5* (Chrna5+, n = 243), only *Syt6* (Syt6+, n = 834), or both *Chrna5* and *Syt6* (Chrna5+Syt6+, n = 564). Expression (copies per million) greater than zero was used as the threshold. 781 cells did not express either *Chrna5* or *Syt6* and were not used in subsequent analyses. Rare cell types with fewer than 10 cells per group are not shown in the heatmap in figure 5 but are included for the differential expression analysis. The FindMarkers function in Seurat was used to identify genes differentially expressed between the Chrna5+ and Chrna5+Syt6+ populations. Adjusted p value < 0.05 was used as the cutoff for identifying differentialy expressed genes. Similar approaches were used to perform the same analyses in multiple cortical regions (Fig S5). L5-6 glutamatergic cells cells in ACA, MOp, SSp, and VISp were taken from the Whole cortex & Hippocampus Smart-Seq (2019) dataset.

### Quantification and Statistical Analysis

Analysis of electrophysiological data was conducted on Clampfit 10.7 and Axograph. Rising slope was measured by fitting a line to the first 50ms of the optogenetic cholinergic responses. Cholinergic response magnitude in voltage clamp was determined by peak current (picoamperes) and charge transfer (picocoulombs) measured by the area under the current response for 1 second.

GCaMP6s imaging data were extracted using the multi-measure feature in Fiji. Maximum projections across time for each experiment were first used to identify acetylcholine-responsive cells and add them to the ROI manager, then fluorescence intensity at all timepoints for each cell was measured. Fluorescence was normalized to the background fluorescence averaged over the first 10 frames. Area under the peak (AUP) of the signal after baseline correction was used to quantify the magnitude of cells’ response to acetylcholine. Percentage response remaining after DHBE and DHBE+Atropine was calculated from the cell’s AUP before and after the blockers. The statistical analysis for electrophysiological recordings and calcium imaging treated each cell as independent with several cells recorded from each slice.

GraphPad Prism 8 was used for statistical analysis and plotting graphs. Bar graphs depicting mean with standard error and boxplots with median and quartiles are shown. Effect sizes are reported as Cohen’s d for major results^96^. Unpaired t-tests or Mann-Whitney tests were used when comparing response properties between cell types, and paired t-test or Wilcoxon test to quantify effect of pharmacological manipulations within cells. Kolmogorov-Smirnov and Fisher’s exact tests were used to compare cumulative distributions and proportions of cells respectively.

## Supplementary Material Legends

**Video S1:** Video shows Chrna5+ neurons in mPFC slices from *Chrna5*-Cre^/+^Ai96^/+^ mice responding to exogenous application of 1mM acetylcholine with an increase in GCaMP6s fluorescence signal

**Video S2:** Video shows Syt6+ neurons from *Syt6*-Cre^/+^Ai96^/+^ mice responding to exogenous application of 1mM acetylcholine with an increase in GCaMP6s fluorescence signal.

**(*Note: Figures S1-5 and Tables S1-3 are in one PDF file. Table S4 is attached as a separate excel file.*)**

**Figure S1.** *Chrna5-Cre* reporter expression in different brain regions.

**Figure S2.** Chrna5+ neurons in primary somatosensory cortex are super-responders to optogenetic acetylcholine release.

**Figure S3.** Chrna5+ cells in primary visual cortex from *Chrna5*-Cre^/+^ Ai14^/+^ Syt6-EGFP^/+^ mice.

**Figure S4**. Chrna5+ acetycholine super-responders in layer 6b.

**Figure S5.** Transcriptomic characterization of Chrna5+ super-responders in other cortical regions.

**Table S1.** Intrinsic electrophysiological properties of Chrna5+ and Chrna5-unlabeled deep-layer neurons in *Chrna5*-Cre^/+^Ai14^/+^ChAT-ChR2^/+^ mice.

**Table S2.** Intrinsic electrophysiological properties of Chrna5+ and Syt6+ deep-layer neurons.

**Table S3.** Comparing expression of major genes modulating postsynaptic cholinergic responses in Chrna5+ and Chrna5+Syt6+ neurons.

**Table S4.** Excel file containing all differentially expressed genes between Chrna5+ and Chrna5+Syt6+ neurons with adjusted p value < 0.05

