## Supplementary Figures S1-S5, supplementary Tables S1-S3 for "Chrna5 and lynx prototoxins identify acetylcholine super-responder subplate neurons"

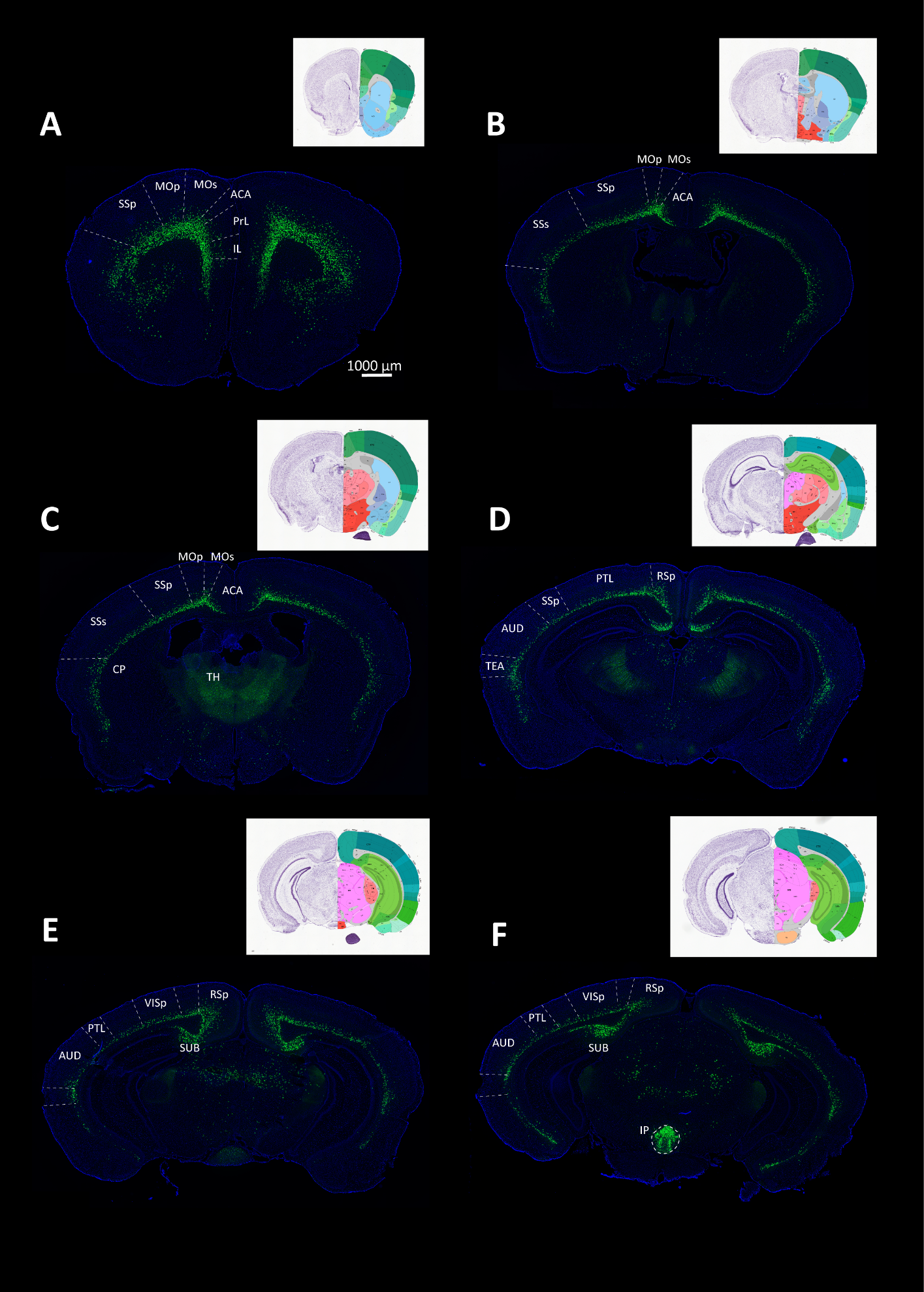

**Figure S1. *Chrna5*-Cre reporter expression in different brain regions.**

Confocal images of coronal brain sections from *Chrna5*-Cre x Ai6 mice expressing the fluorescent reporter Zsgreen in Chrna5+ cells are shown. Reference images in the inset are from the Allen brain reference atlas. Chrna5+ cells are found in layer 6 across the cortical mantle: **A,** Prelimbic (PrL), infralimbic (IL), anterior cingulate area (ACA) (related to Fig 1 in main manuscript), **B-C**, primary motor (MOp), and primary somatosensory (SSp) areas. **D-E**, Chrna5+ cells are also found in layer 6 of posterior cortical areas including parietal (PTL), retrosplenial (RSp), Auditory (AUD) and primary visual (VISp) cortex. **F**, Subcortically, Chrna5+ cells are found in the Subiculum (SUB) and the interpeduncular (IP) nucleus as previously reported in ^1^.

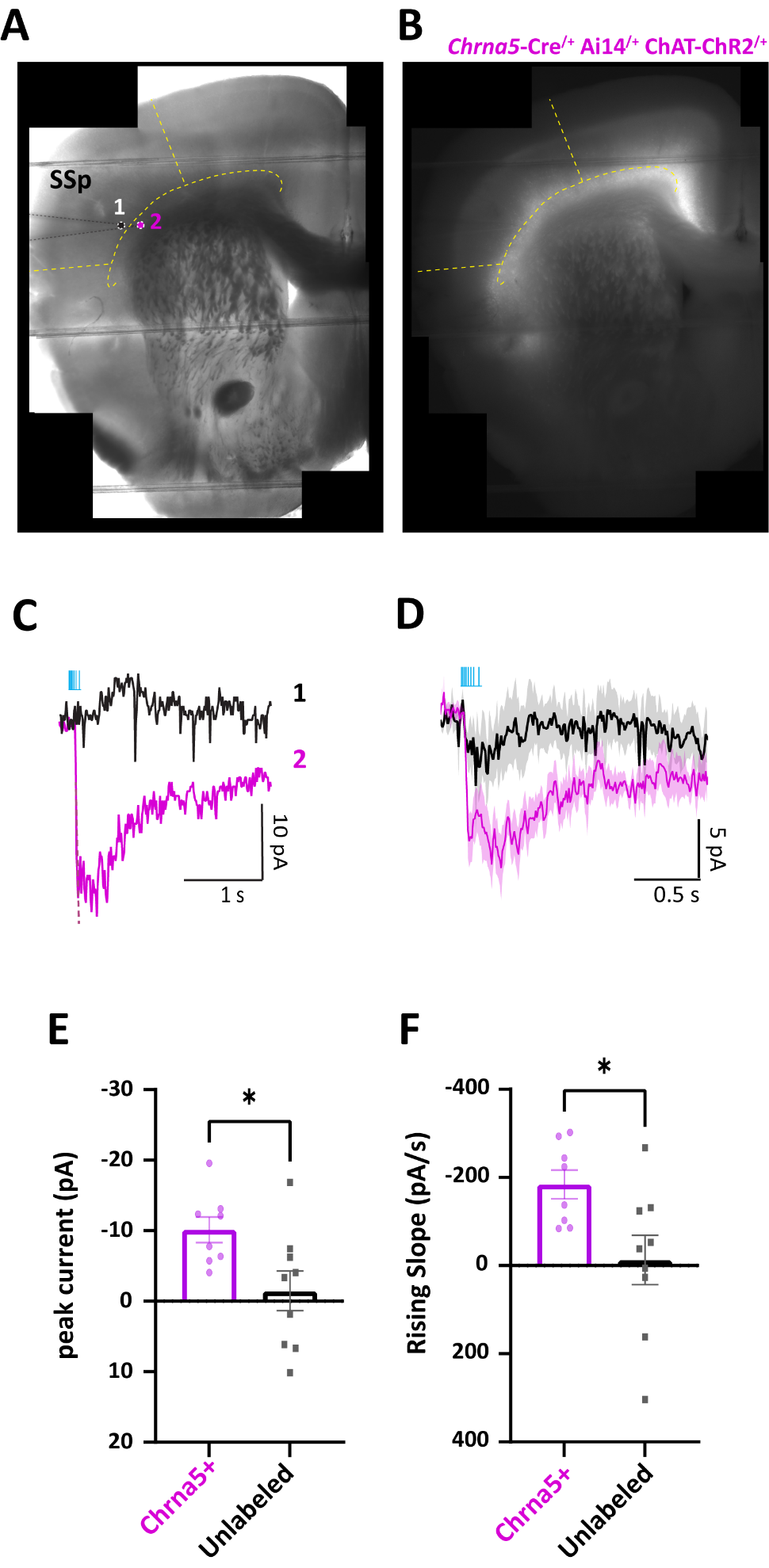

**Figure S2. Chrna5+ neurons in primary somatosensory cortex are super-responders to optogenetic acetylcholine release.**

Slice patch-clamp electrophysiological recordings in primary somatosensory cortex (SSp) of *Chrna5*-Cre^/+^ ChAT-ChR2^/+^ mice (related to Fig 1 in main manuscript). **A,** IRDIC and **B,** tdTomato fluorescence image of coronal brain slice. The position of a recorded unlabeled neuron is indicated by black circle (neuron 1). The patch pipette is positioned on a *Chrna5*-labeled cell (neuron 2, pink circle) in layer 6b. The boundary of layer 6b is indicated by yellow dotted lines. Note, the horizontal lines are from the harp holding the slice down. **C**, Opto-cholinergic responses from unlabeled neuron 1 and Chrna5+ neuron 2 are shown. Chrna5+ neuron in layer 6b has a strong and fast nicotinic current in response to endogenous acetylcholine release unlike unlabeled neuron 1, which has a slow outward current. **D,** Average light-evoked cholinergic responses from Chrna5+ versus unlabeled neurons are shown. Chrna5+ cells in SSp have stronger and faster responses. **E-F,** Bar graphs quantifying the peak current (E) and rising slope (F) of optogenetic cholinergic responses in Chrna5+ and unlabeled cells. **P* < 0.05, unpaired t-test.

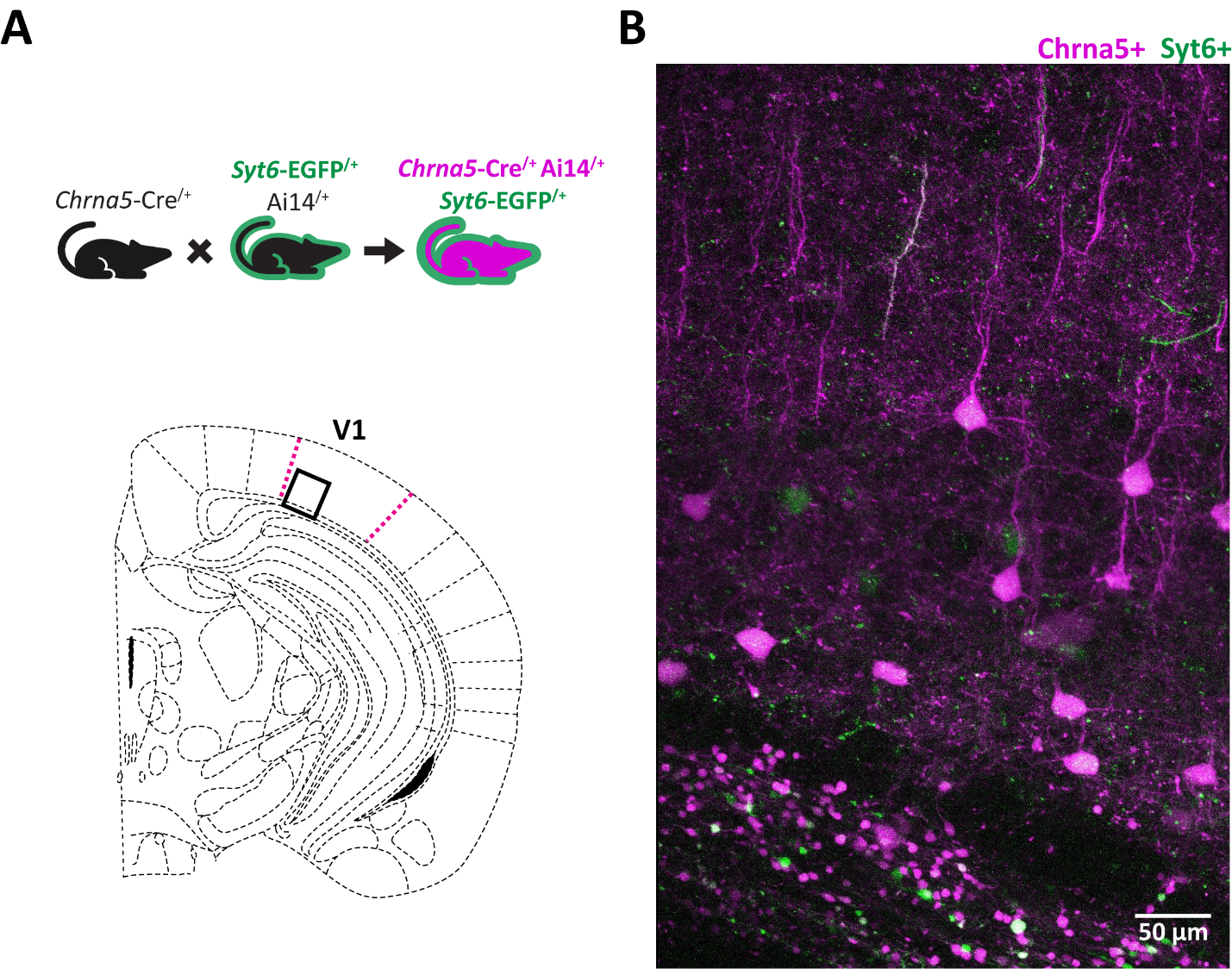

**Figure S3. Chrna5+ cells in primary visual cortex from *Chrna5*-Cre^/+^ Ai14^/+^ Syt6-EGFP^/+^ mice.**

**A,** Schematic showing generation of double labeled *Chrna5*-Cre^/+^ Ai14^/+^ Syt6-EGFP^/+^ mice (top) and location of the region of interest (bottom, ^2^) for two-photon imaging in primary visual cortex. (Related to Fig 1G-I in main manuscript). **B**, *Chrna5*-Cre tdTomato labeled neurons are found in layer 6 and layer 6b of primary visual cortex, while *Syt6*-EGFP expression is greatly reduced consistent with previous reports in GENSAT ^3,4^

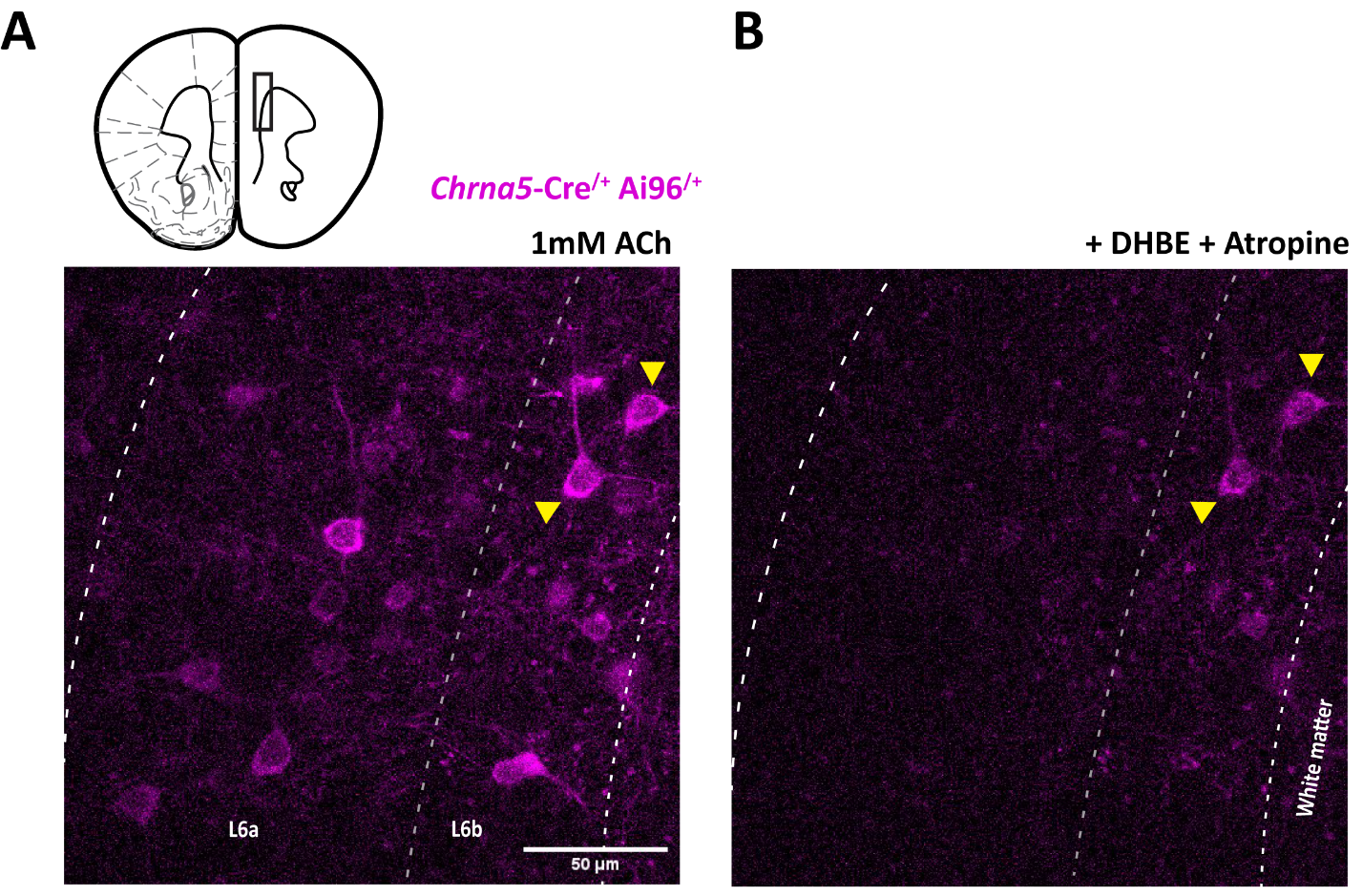

**Figure S4. Chrna5+ acetycholine super-responders are in layer 6b.**

**A,** Two photon Ca^2+^ imaging in a prefrontal brain slices from *Chrna5*-Cre^/+^Ai96^/+^ mice showing acetylcholine-evoked GCaMP6s responses in Chrna5+ neurons (Related to Fig 2 in manuscript). **B**, Location of Chrna5+ neurons resilient to competitive nicotinic antagonist DHBE + atropine is indicated by yellow arrows. These acetylcholine super-responder neurons are restricted to layer 6b.

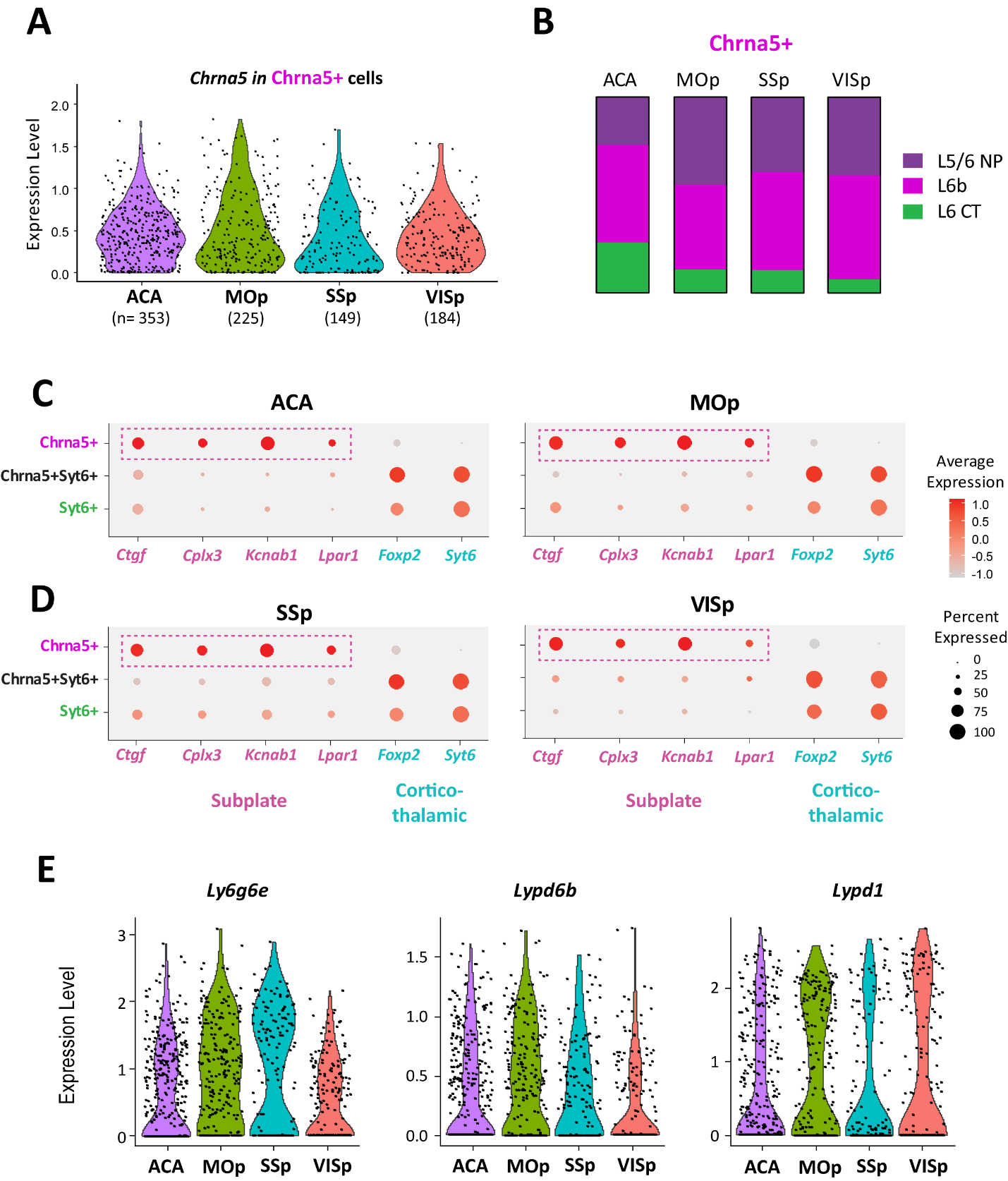

**Figure S5. Transcriptomic characterization of Chrna5+ super-responders in other cortical regions.**

**A,** Violin plots show expression level of *Chrna5* in Chrna5+ neurons across anterior cingulate (ACA), primary motor (Mop), somatosensory (SSp), and visual (VISp) cortices. **B,** Bars indicate proportions of Chrna5+ neurons in ACA, Mop, SSp, and VISp that are L5/6NP (Near-projecting), L6b, or L6CT (Corticothalamic). These 3 cell subclasses together represent >80% of Chrna5+ neurons with L6b and L5/6NP constituting the majority. **C-D,** Dotplots show the expression of subplate and corticothalamic marker genes in ACA, MOp (C), and SSp, VISp (D). Dot size indicates the percentage of cells within each group expressing that gene, color of the dot indicates average expression level relative to other groups. Dotted box indicates the selective enrichment of subplate marker genes in Chrna5+ neurons across cortical regions. **E**, Violin plots show expression of Lynx genes *Ly6g6e*, *Lypd6b* and *Lypd1* known to modulate nicotinic receptor properties in Chrna5+ neurons in ACA, MOp, SSp, and VISp. (See Fig 3 in main manuscript describing similar findings in ACA). Data from the Allen Institute ^5^

**Table S1**

|  | **Chrna5+**  (n = 24) | **Chrna5-**  (n = 13) | **Unpaired t-test** |
| --- | --- | --- | --- |
| Resting membrane potential (mV) | -85 ± 1 | -85 ± 1 | t_(35)_ = 0.63, *P* = 0.53 |
| Input resistance (MΩ) | 133 ± 12 | 167 ± 18 | t_(35)_ = 1.68, *P* = 0.10 |
| Membrane capacitance (pF) | 54 ± 2 | 57 ± 2 | t_(35)_ = 1.05, *P* = 0.30 |
| Spike threshold (mV) | -49 ± 1 | -49 ± 1 | t_(35)_ = 0.34, *P* = 0.73 |
| Spike amplitude (mV) | 74 ± 2 | 73 ± 3 | t_(35)_ = 0.21, *P* = 0.83 |
| Rheobase (pA) | 135 ± 11 | 101 ± 11 | t_(33)_ = 1.92, *P* = 0.06 |

**Intrinsic electrophysiological properties of Chrna5+ and Chrna5- unlabeled deep-layer neurons in *Chrna5*-Cre^/+^Ai14^/+^ChAT-ChR2^/+^ mice.** None of the intrinsic properties are significantly different between the Chrna5+ and unlabeled Chrna5- neurons in Figure 1.

**Table S2**

|  | **Chrna5+**  (n = 12) | **Syt6+**  (n = 14) | **Unpaired t-test** |
| --- | --- | --- | --- |
| Resting membrane potential (mV) | -86 ± 1 | -88 ± 1 | t_(24)_ = 1.65, *P* = 0.12 |
| Input resistance (MΩ) | 156 ± 17 | 123 ± 6 | t_(24)_ = 1.95, *P* = 0.06 |
| Membrane capacitance (pF) | 55 ± 3 | 59 ± 2 | t_(24)_ = 1.02, *P* = 0.32 |
| Spike threshold (mV) | -50± 1 | -49 ± 1 | t_(24)_ = 0.61, *P* = 0.55 |
| Spike amplitude (mV) | 79 ± 2 | 73 ± 3 | t_(24)_ = 1.59, *P* = 0.12 |
| Rheobase (pA) | 117 ± 21 | 126 ± 13 | t_(22)_ = 0.37, *P* = 0.71 |

**Intrinsic electrophysiological properties of Chrna5+ and Syt6+ deep-layer neurons.** None of the intrinsic properties are significantly different between Chrna5+ and Syt6+ neurons in Figure 3.

**Table S3**

| Genes | P value | Chrna5+  cells % | Chrna5+Syt6+  cells % | Adjusted P value | Fold change  (Chrna5+/ Chrna5+Syt6) |
| --- | --- | --- | --- | --- | --- |
| ***Lypd1*** | 3.40E-23 | 0.593 | 0.381 | 9.88E-19 | **2.55** |
| ***Ly6g6e*** | 2.13E-25 | 0.671 | 0.434 | 6.20E-21 | **2.03** |
| ***Lypd6b*** | 8.81E-45 | 0.539 | 0.143 | 2.56E-40 | **1.51** |
| *Ache* | 1.43E-23 | 0.889 | 0.784 | 4.15E-19 | 1.50 |
| *Chrm2* | 9.06E-38 | 0.428 | 0.086 | 2.63E-33 | 1.25 |
| *Ly6h* | 7.69E-07 | 1 | 1 | 0.02 | 1.10 |
| *Lypd6* | 5.95E-19 | 0.136 | 0.011 | 1.73E-14 | 1.07 |
| *Chrna4* | 0.089 | 0.967 | 0.978 | 1 | 1.05 |
| *Chrnb2* | 0.079 | 0.44 | 0.374 | 1 | 1.02 |
| *Chrnb3* | 0.045 | 0.132 | 0.088 | 1 | 1.02 |
| *Chrna7* | 0.139 | 0.132 | 0.1 | 1 | 1.01 |
| *Chrm4* | 0.839 | 0.156 | 0.156 | 1 | 1.01 |
| *Chrna2* | 0.351 | 0.008 | 0.004 | 1 | 1.00 |
| *Chrnb4* | 0.591 | 0 | 0.001 | 1 | 1.00 |
| *Chrna3* | 0.088 | 0.235 | 0.295 | 1 | 1.00 |
| *Chrm3* | 0.025 | 0.687 | 0.896 | 1 | 0.99 |
| *Lynx1* | 1.39E-05 | 0.918 | 0.953 | 0.40 | 0.91 |
| *Chrm1* | 1.69E-06 | 0.827 | 0.902 | 0.05 | 0.89 |
| *Ly6e* | 3.35E-10 | 1 | 1 | 9.72E-06 | 0.88 |
| *Chrna5* | 5.38E-13 | 1 | 1 | 1.56E-08 | 0.80 |

**Comparing expression of major genes modulating postsynaptic cholinergic responses in Chrna5+ and Chrna5+Syt6+ neurons.** Genes of interest for cholinergic properties filtered from the list of all genes in our differential expression analysis. Genes are sorted in descending order of the fold change between Chrna5+ and Chrna5+Syt6+ neurons. 3 Lynx prototoxins- Lypd1, Ly6g6e, and Lypd6b (bold text) are the highest enriched among these cholinergic modulatory genes in Chrna5+ neurons (Related to Fig 3 in manuscript).
